# Roles of the MO25 protein Pmo25 in contractile-ring stability and localization of the NDR kinase Sid2 during cytokinesis

**DOI:** 10.1101/2025.05.13.653815

**Authors:** Yanfang Ye, Sha Zhang, Jack R. Gregory, Aysha H. Osmani, Evelyn G. Goodyear, Jian-Qiu Wu

## Abstract

Mouse protein-25 (MO25) family proteins are crucial in development and morphogenesis from plants to humans. The fission yeast MO25 protein Pmo25 is essential for cell polarity and division. However, how Pmo25 regulates cytokinesis remains largely unknown. Here we found that the actomyosin contractile ring and septum formation were defective during cytokinesis in *pmo25* mutants. Pmo25 physically and genetically interacted with the myosin-II light chain Cdc4, which is essential for the contractile-ring assembly and function. Additionally, *pmo25* mutations had synthetic genetic interactions with all other tested mutations in contractile-ring proteins. Moreover, Pmo25 colocalized with the NDR kinase Sid2 and participated in its recruitment to the division plane. Furthermore, Pmo25 directly bound the Munc13/UNC-13 protein Ync13 and modulated the secretion of glucanase Eng1 to the division site for daughter-cell separation. Our data provide insight into how Pmo25 regulates cytokinesis and suggest that the conserved MO25 proteins can link various steps of cytokinesis.

## INTRODUCTION

Cell division and cell morphogenesis are fundamental determinants of cell proliferation, differentiation, and development. The cell cycle transitions must be highly regulated to ensure that these processes occur properly. This raises the question of how cells maintain the dynamic transition of interconnected signaling networks and signal fidelity. The mechanisms and signaling cascades that control cell growth and division are highly conserved in various aspects between yeasts and higher eukaryotes^1-7^. The rod-shaped fission yeast *Schizosaccharomyces pombe* grow from cell ends and divide by medial fission. Thus, it is an ideal, genetically tractable model system to elucidate these conserved mechanisms and signaling pathways.

Cytokinesis is the last step of the cell-division cycle, which partitions cytoplasm and organelles from a mother cell to two daughter cells. In most eukaryotic cells from fungi to mammalian cells, cytokinesis occurs with several crucial steps: cleavage site selection; actomyosin contractile-ring assembly, maturation, constriction; plasma membrane deposition at the division plane; and extracellular matrix formation or remodeling^8-12^. Cytokinesis also needs the cooperation of signaling pathways and membrane trafficking, including exocytosis and endocytosis, to succeed^13-18^. The Septation Initiation Network (SIN) controls growth at the division site to guide cytokinesis in *S. pombe*. The SIN pathway resembles the Mitotic Exit Network in budding yeast and the mammalian Hippo pathways^3,5,19-21^. It is essential for contractile-ring assembly, maintenance, and constriction, as well as for the formation of the septum and new cell ends^22-24^. The signaling of the SIN pathway is controlled by the GTPase Spg1, which in turn activates three downstream kinases: the Hippo-like kinase Cdc7, the PAK-related GC kinase Sid1, and the NDR kinase Sid2. Apart from Sid2 with its regulatory subunit and activator Mob1, most SIN components are situated at the spindle pole body (SPB)^25,26^, which is functionally equivalent to centrosomes. Upon activation, the Sid2-Mob1 complex translocates from the SPBs to the medial division site where it facilitates contractile-ring assembly and constriction as well as septum formation^27^. Errors in SIN signaling cause cytokinesis failure. However, cells continue their growth and nuclear cycle, resulting in elongated and multinucleated cells before cell lysis. It has been reported that the SIN cascade inhibits the Morphogenesis Orb6 Network (MOR) signaling pathway in mitosis by interfering with the activation of the most downstream MOR component, Orb6, which is the other NDR kinase in fission yeast^1,28,29^. The MOR signaling regulates actin assembly and is essential for cell polarity and morphogenesis^30-33^. In addition, the SIN kinases Cdc7 and Sid1 also regulate the MOR components, such as the localization of the MO25 protein Pmo25 at the SPBs and the kinase activity of Orb6 during interphase^33,34^.

MO25 was originally identified as a gene expressed at the early cleavage stage of mouse embryogenesis^35^. It is a scaffold protein that forms a heterotrimeric complex with STRAD and the tumor suppressor liver kinase 1 (LKB1). The LKB1-STRAD-MO25 heterotrimeric complex stabilizes a closed conformation of STRAD and triggers LKB1 nucleocytoplasmic shuttling and activation^36-39^. LKB1 modulates several cellular processes, such as cell growth and polarity^40,41^. Besides its role in the heterotrimeric complex, MO25 is also a key regulator of ion homeostasis and development/morphogenesis^42^. MO25 proteins are highly conserved in eukaryotic cells from plants to humans^43-46^. In budding yeast, the MO25-like protein Hym1 functions in the RAM network, which is equivalent to the MOR pathway in *S. pombe* and is essential to cell polarity, morphogenesis, and daughter-cell separation^4,47,48,49,50^. In addition, Hym1 is involved in the G1 to S phase transition^51^. In *Neurospora crassa*, HYM1/MO25 controls the NDR kinase module as well as MAP kinase signaling during intercellular communication^51^. Similarly, Pmo25, a component of the MOR network in fission yeast, functions as an upstream regulator of the MOR network and mediates signaling connection between the SIN and MOR^33,34,52^. Pmo25 has also been reported to interact with the GC kinase Ppk11^53^. *pmo25* mutants are defective in cell morphogenesis and daughter-cell separation^33,34,52^. However, unlike its roles in cell morphogenesis, how Pmo25 regulates cell separation and its binding partners during cytokinesis remains largely unknown.

The fission yeast Ync13 is a member of the Munc13/UNC-13 protein family, which is conserved in plants, fungi, and humans^54-58^. Ync13 is essential for cytokinesis in rich media, and its mutants are defective in septum formation, resulting in cell lysis during daughter-cell separation ^58^. The Munc13/UNC-13 proteins have conserved functions in vesicle priming and tethering before vesicle fusion with the plasma membrane during exocytosis^54-57,59-65^. Ync13 modulates the recruitment, maintenance, and spatial distribution of cell wall enzymes including the glucan synthases Bgs4 and Ags1, and the glucanase Eng1 during cytokinesis^58,66-77^. However, the function of Ync13 in cytokinesis remains poorly understood.

In this study, we demonstrate that Pmo25 is not only required for cell separation in late cytokinesis but also for contractile-ring stability. Pmo25 interacts with the myosin-II essential light chain Cdc4 and the NDR kinase Sid2. Pmo25 is also involved in recruiting Sid2 to the division plane. In addition, we reveal that Pmo25 directly binds Ync13 and provide evidence that Pmo25 regulates Ync13 and Eng1 levels at the division site. Collectively, our data suggest that Pmo25 interacts with several cytokinetic proteins and links various steps of cytokinesis.

## RESULTS

### Pmo25 is important for contractile-ring formation, stability, and septum formation during cytokinesis

Despite previous studies of Pmo25^33,34,52^, its role in cytokinesis remains poorly understood. As Pmo25 is an essential gene, to further explore the functions of Pmo25 in cytokinesis, we created new mutants of *pmo25* using the marker reconstitution mutagenesis method by replacement of the chromosomal *pmo25* gene with error-prone PCR products amplified using full length *pmo25^+^* as previously described^78-81^. *pmo25-21* showed similar polarity and septation defects as other previously characterized mutants *pmo25-2* and *pmo25-35*^33,34^ at 36°C (Figures 1A-1C). Cells gradually lost polarity and became rounder with extended growth at 36°C (Figure 1C). Even at 25°C, *pmo25-21* cells experienced delayed septation. We used the plasma membrane-associated serine-rich cell wall sensor Wsc1 to examine the plasma membrane, α-tubulin Atb2 or the SPB protein Sad1 to examine the cell cycle stages, and the myosin-II regulatory light chain Rlc1 or the F-BAR protein Cdc15 to examine the contractile ring during cytokinesis in WT and *pmo25* mutants (Figure 1). In WT cells, the contractile ring assembled from the precursor nodes, dots-shaped protein complexes on the plasma membrane at the division site, and constricted to guide plasma membrane invagination and septum formation (Figures 1A and 1B) as reported^9,82,83^. In *pmo25-21* mutant at 25°C, Rlc1-mCherry nodes appeared at the division site ∼10.5 min before SPB separation, and the compact-ring formation (from nodes appearance to condensation into a ring without lagging nodes), maturation (from a compact ring to the start of fast phase of ring constriction^84^), constriction (from the start of fast-phase of ring constriction^84^ to the ring constricted to a spot at cell center with highest Rlc1 pixel intensity), disassembly (from ring spot to Rlc1 disappearance from the division site), and septum maturation (from Rlc1 disappearance to the start of daughter-cell separation) took ∼29.2 min, ∼7.4 min, ∼37.7 min, ∼24.0 min, and >72.5 min (only quantified the separated cells by the end of the movie so the septum maturation time was underestimated), respectively. All these processes showed significant delay compared to WT cells that took ∼24.7 min, ∼5.1 min, ∼27.2 min, ∼10.1 min, and ∼16.2 min (p < 0.01), respectively (Figures 1A and 1B). Thus, suggesting that Pmo25 plays multiple important roles in cytokinesis.

**Figure 1.**
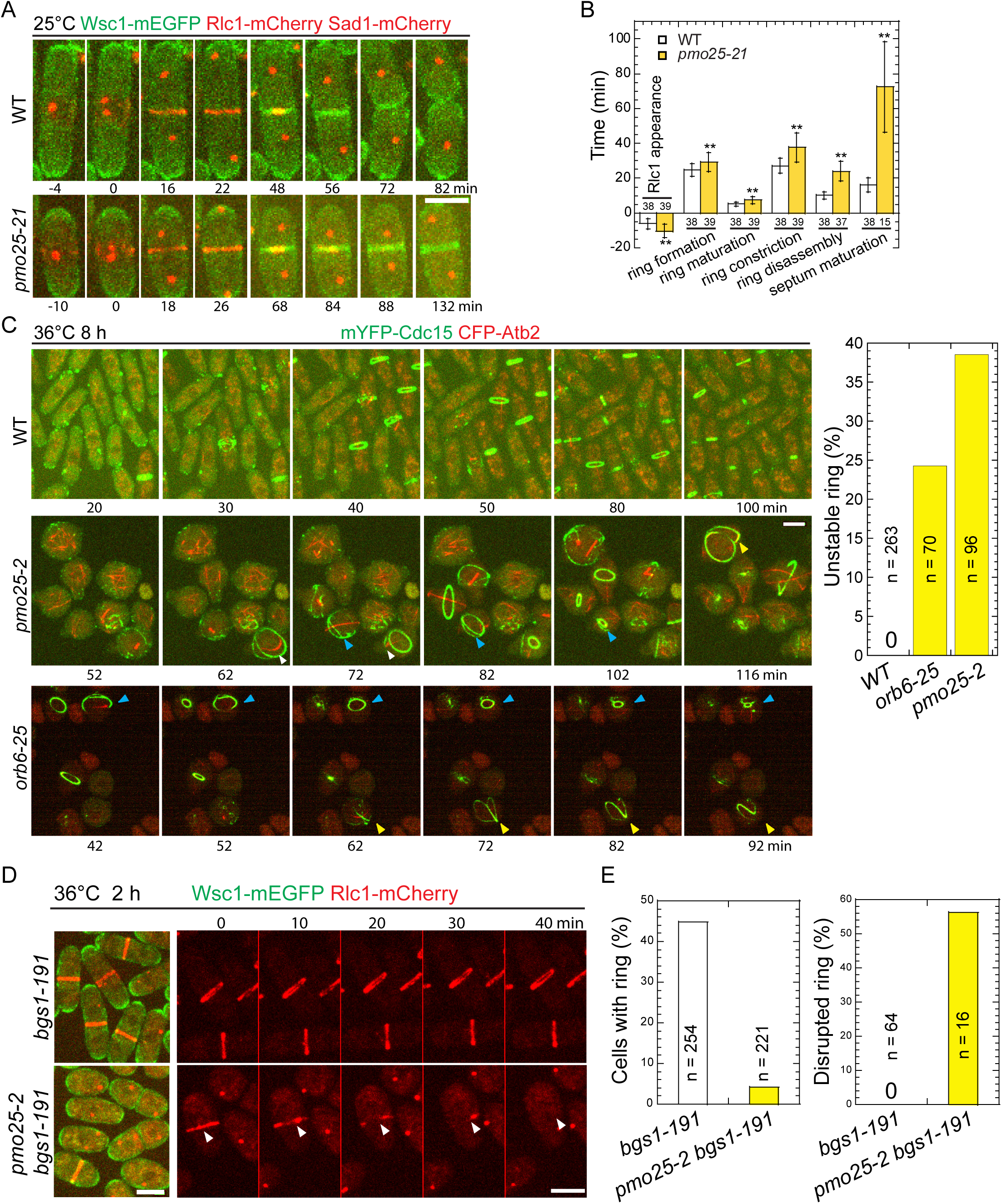
pmo25 mutants are defective in cytokinesis. (A and B) Time course (A) and quantification (B) of the main cytokinesis events (in min) of cells expressing Wsc1-mEGFP Rlc1-mCherry Sad1-mCherry in WT (JW8614) and *pmo25-21* mutant (JW8615) at 25°C. Time <vol>0 marks the separation of two SPBs. Numbers of cells analyzed are shown above or below the bars. \*\**p* < 0.001. (C) Time lapse (in min) of mYFP-Cdc15 CFP-Atb2 in WT (JW3186), *pmo25-2* (JW8148), and *orb6-25* (JW8333) cells. Cells were grown in log phase at 25°C for ∼36 h, then shifted to 36°C for 8 h before imaging at ∼36°C. Times are minutes after the start point of imaging. Yellow, white, and blue arrowheads indicate the same cells that had an abnormal ring during the movie. Right, quantification of unstable contractile rings (the rings became frayed, deformed, or disintegrated before the completion of its constriction) in WT, *orb6-25*, and *pmo25-2* cells during the 2 h movies at 36°C after cells were shifted to 36°C for 8 h. (D and E) Micrographs (D) and quantification (E) of Wsc1-mEGFP Rlc1-mCherry in *bgs1-191* (JW2766) and *pmo25-2 bgs1-191* mutant (JW8575) at 36°C. Cells were grown in log phase at 25°C for ∼36 h, then shifted to 36°C for 2 h before imaging at 36°C. (D) Left, single time-point image; Right, time lapse (in min) shown Rlc1 signal only. Time 0 marks the start of the movie. Arrowheads mark a collapsed ring. (E) Left and right graphs show percentage of cells with contractile ring and ring break-down before fully constriction during the time-lapse movies, respectively. Bars, 5 μm.

When grown longer at 36°C, unlike in WT, 39% of the contractile ring became frayed, deformed, or disintegrated before or during its constriction in *pmo25-2* mutant (Figure 1C). These defects could not be explained solely by the round shape of *pmo25-2* cells because the ring defects in polarity mutant *orb6-25* were less frequent and less severe (Figure 1C and Movie 1). Based on these results, we hypothesized that Pmo25 may also affect the contractile ring integrity or stability. To test this hypothesis further, we examined the ring stability in the (1,3)β-glucan synthase mutant *bgs1/cps1-191*, which significantly delays contractile-ring constriction with a stable contractile ring^74,75,85^. As expected, the percentage of *bgs1-191* cells with rings reduced dramatically by the *pmo25* mutation. After shifting to 36°C for 2 h, ∼45% of *bgs1-191* cells had a contractile ring, whereas only ∼5% of *bgs1-191 pmo25-2* cells had a ring (Figures 1D and 1E). We reasoned that the lack of the ring in *bgs1-191 pmo25-2* cells may have been caused by faster constriction and/or ring collapse. In 70-min time lapse movies at 36°C, essentially all the rings in *bgs1-191* cells remained stable and did not constrict obviously. In contrast, ∼55% of the rings in *bgs1-191 pmo25-2* cells were disrupted or collapsed during the movie (Figures 1D and 1E, Movies 2 and 3). These results suggested that Pmo25 is important to contractile-ring stability/integrity during cytokinesis.

Temperature-sensitive *pmo25* mutants may still retain some Pmo25 functions even at 36°C. To confirm Pmo25’s role in contractile-ring stability, we examined cytokinesis in the *pmo25*Δ (Δ means deletion) mutant grown at 25°C using the Tetrad Fluorescence Microscopy, which facilitates the observation of terminal phenotype of essential genes^86-88^. As reported^33,34^, *pmo25*Δ cells became rounder, ceased growing, and lysed after several divisions, in contrast to the rod-shaped WT *pmo25*^+^ cells (*hph^S^*) after tetrad dissection of *pmo25*^+^/*pmo25*Δ heterozygous diploid cells (Figure 2A). Similar to the *pmo25* temperature-sensitive mutants, *pmo25*Δ cells exhibited a delay in cytokinesis. Except for the ring sliding from the cell center, ∼25% of contractile rings in *pmo25*Δ, but not in WT cells, collapsed during formation, maturation, or constriction (Figures 2B and 2C, Movies 4 and 5). Although the ring constriction took longer in *pmo25*Δ cells, ∼52.3 min compared to ∼27.8 min in WT cells (Figure 2D; p < 0.001), the constriction rate showed no obvious difference, ∼0.39 μm/min in *pmo25*Δ cells compared to ∼0.41 μm/min in WT cells (p = 0.51). These results indicate that Pmo25 is important for contractile-ring stability and integrity during cytokinesis.

**Figure 2.**
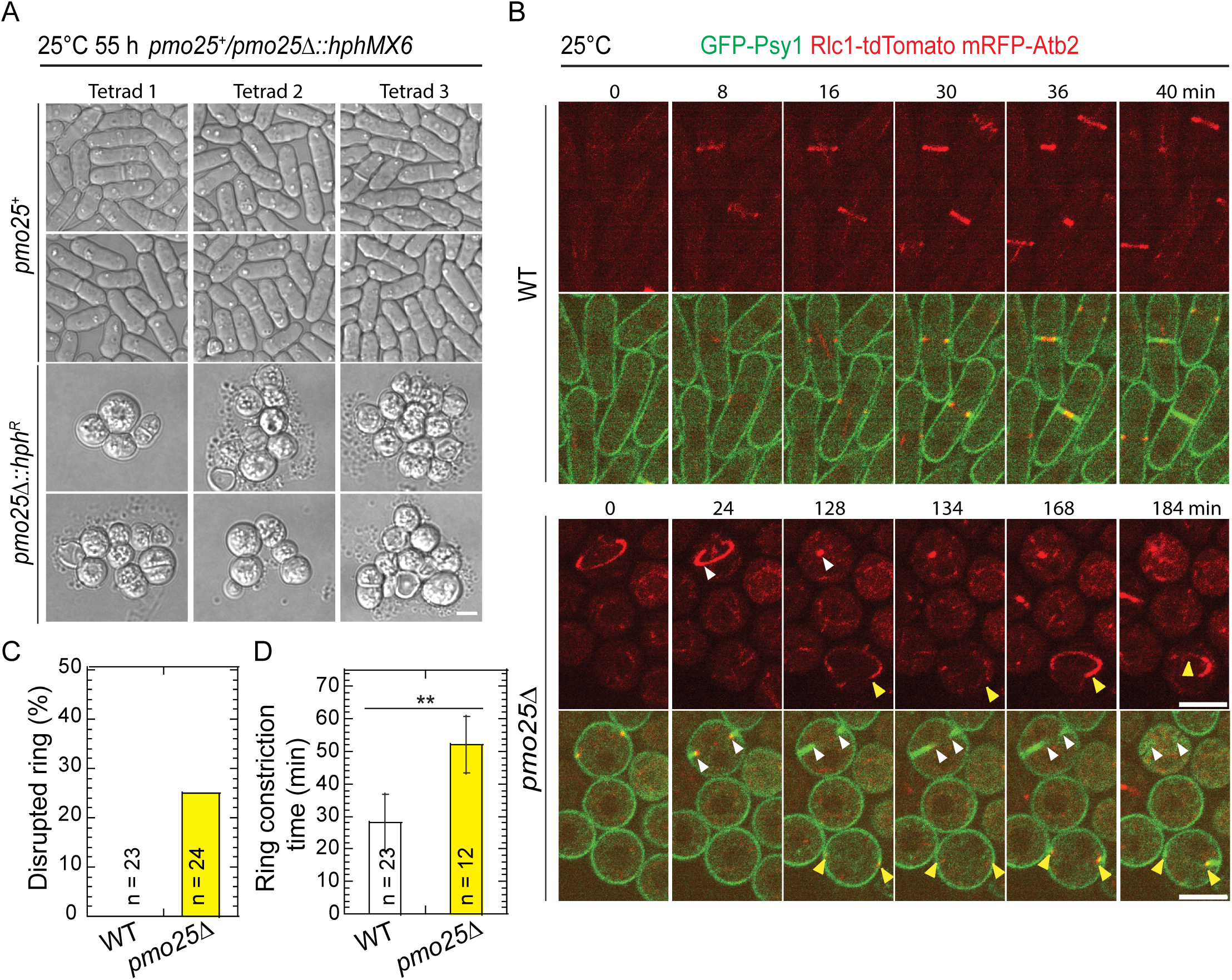
Pmo25 is important for cell polarization and contractile-ring stability. (A) Morphology of *pmo25Δ* mutant. One copy of *pmo25* was deleted in diploid cells. After tetrad dissection, spores were grown at 25°C for ∼55 h, then imaged under microscope and tested for hygromycin resistance (*hph^R^, pmo25Δ*) or sensitivity (*hph^S^, pmo25^+^*). (B) Time courses (in min) of GFP-Psy1 Rlc1-tdTomato RFP-Atb2 in WT and *pmo25Δ* cells at 25°C. Time 0 is the start of the movies. Upper panel, Rlc1 and Atb2 signals; Lower panel, merge. Arrowheads mark examples of disrupted rings. Bars, 5 μm. (C) Percentage of cells with contractile ring break-down before or during constriction in movies (≥3 hrs) as in (B). (D) Times of the contractile ring constriction in WT and *pmo25Δ* cells measured from movies as in (B). \*\**p* < 0.001.

### Pmo25 binds Cdc4 and is important for Cdc4 ring stability

Cells lacking Pmo25 lead to contractile-ring instability during its formation or constriction, suggesting that Pmo25 may interact with other proteins in the contractile ring. To test this idea, we first tested if Pmo25 interacts with the essential contractile ring components such as the IQGAP scaffold protein Rng2, the formin Cdc12, the F-BAR protein Cdc15, the myosin II heavy chain Myo2, and the essential light chain Cdc4 by yeast-two-hybrid assays. Although not very strong, a positive interaction between Pmo25 and Cdc4 was detected (Figure 3A). To address whether Pmo25 binds Cdc4 in fission yeast cells, we performed Co-IP and found that Pmo25-13Myc co-immunoprecipitated with mYFP-Cdc4 (Figure 3B).

**Figure 3.**
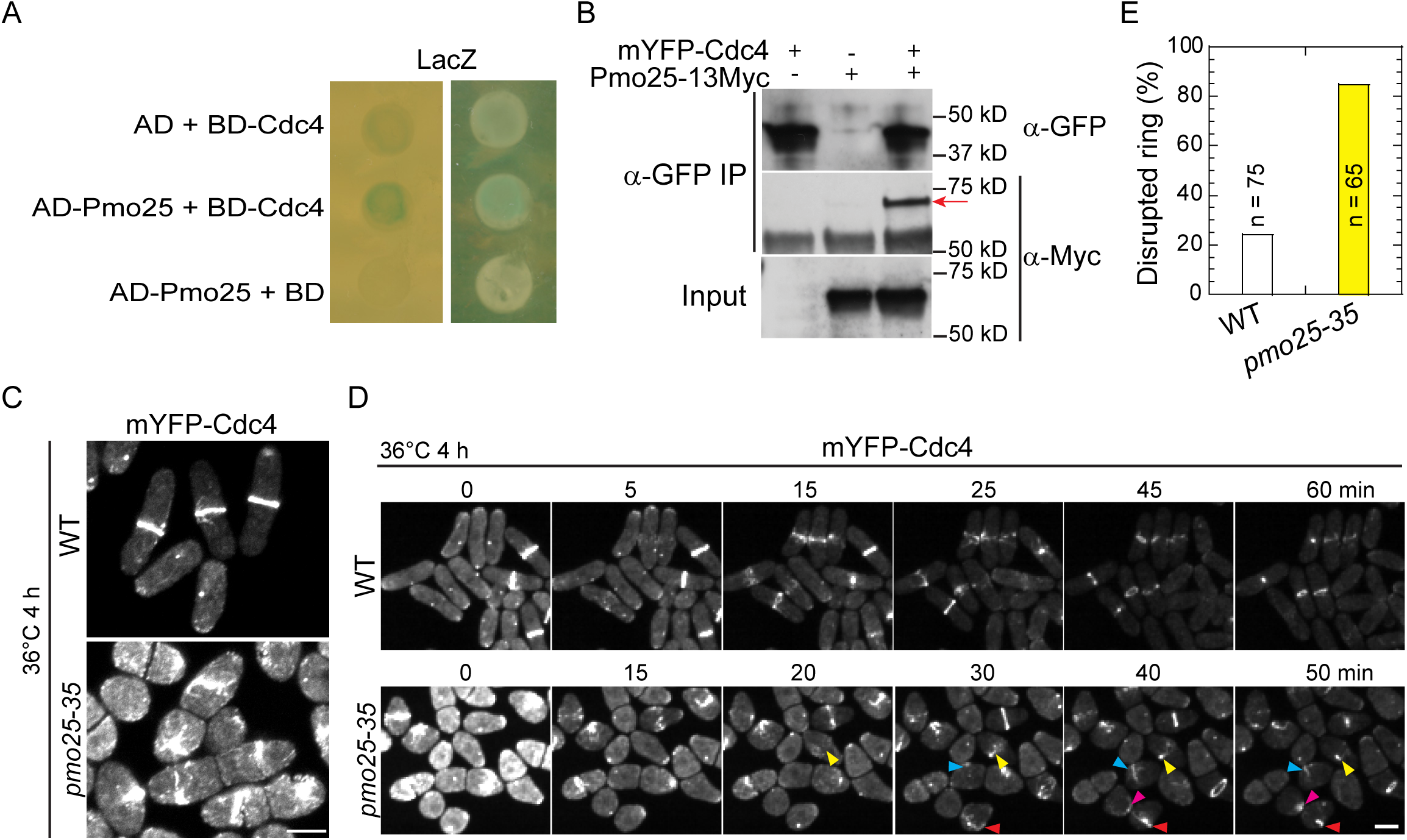
Pmo25 interacts with the myosin-II essential light chain Cdc4. (A) Pmo25 and Cdc4 interact in yeast two-hybrid assays. *S. cerevisiae* MaV203 cells were transformed with AD-Pmo25 and/or BD-Cdc4 with LacZ as a reporter. (B) Co-IP of Pmo25 and Cdc4. Extracts of cells expressing Pmo25-13Myc and/or mYFP-Cdc4 (JW910, JW8662 and JW9877) were immunoprecipitated with anti-GFP polyclonal antibody, then detected by anti-Myc or anti-GFP monoclonal antibodies, respectively. The arrow marks the Pmo25-13Myc band. (C-E) *pmo25* affects mYFP-Cdc4 ring stability. Single image (C) and selected images from time lapse (D, in min) of mYFP-Cdc4 in WT (JW10097) and *pmo25-35* mutant (JW10098) grown at 36°C for 4 h before imaging at 36°C. Arrowheads of the same color mark the same cell. (E) Quantification of cells with mYFP-Cdc4 ring break-down before or during ring constriction in movies as in (D). Bars, 5 μm.

To dissect the contribution of Pmo25 to Cdc4 function, we examined Cdc4 localization in *pmo25* mutants. Neither COOH- nor NH_2_-tagged Cdc4 is fully functional; therefore, we used mYFP-Cdc4, which is more functional, to observe Cdc4 localization at the division site in *pmo25* mutant cells. After shifting the cells to 36°C for 4 h, ∼20% of mYFP-Cdc4 rings collapsed in *pmo25^+^* cells; in contrast, >80% of mYFP-Cdc4 rings collapsed and disrupted in *pmo25-35* cells (Figures 3C-3E). In addition, we observed strong synthetic genetic interactions between *pmo25* mutations and all other tested mutations in the core proteins of the contractile ring, such as the myosin-II light chains *rlc1* and *cdc4*, the formin *cdc12*, the IQGAP *rng2*, and the myosin-II heavy chains *myo2* (Table 1 and Figure S1). Incomplete and disorganized septa were more frequently observed in the multinucleated double mutants than in the single mutants. These results indicate that Pmo25 associates with Cdc4 and plays a role in contractile-ring stability during cytokinesis.

**TABLE 1:**
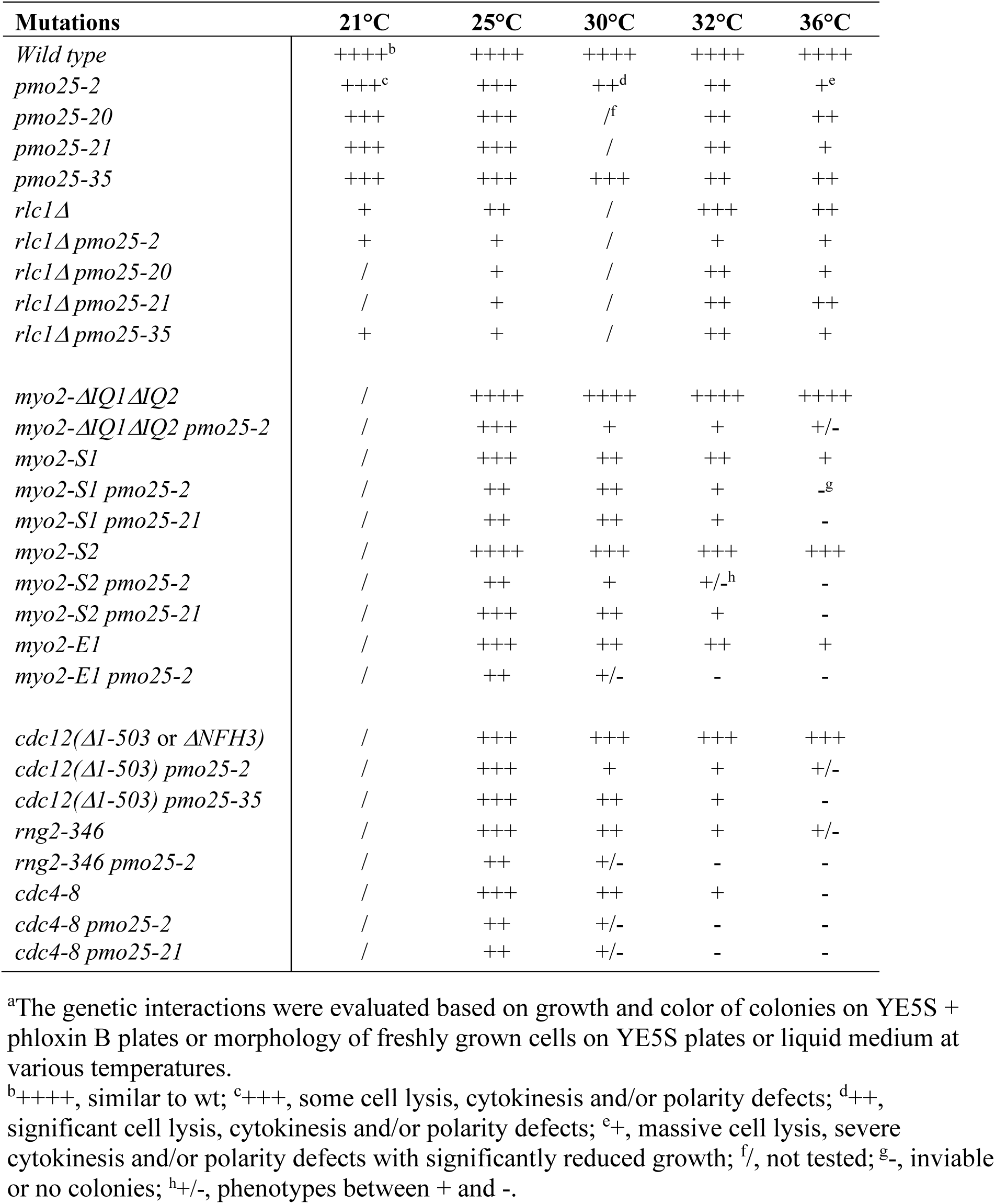
Genetic interactions between mutations in pmo25 and genes in the contractile ring^a^.

### Pmo25 is involved in the recruitment or maintenance of the SIN protein complex Sid2/Mob1 at the division site

Pmo25 mediates the signaling linkage between the SIN and the MOR network for cell morphogenesis/separation following cytokinesis^33,53^. Pmo25 localizes mainly at the SPB that has the active SIN signaling, which is controlled by the Cdc7 and Sid1 kinases but not by the Sid2 kinase^33,34^. Inactivation of the SIN signaling leads to a failure of contractile-ring maintenance, constriction, and septum formation, ultimately leading to cell lysis during cytokinesis due to the defective septum^8,12,24,89^. The cytokinetic defects observed in *pmo25* mutants prompted us to investigate if Pmo25 may feedback to the SIN signaling. We first tested Cdc7 localization in *pmo25* mutant cells 4 h after shifting cells to 36°C. In WT cells, consistent with previous reports^90-93^, Cdc7-YFP localized to the two SPBs at early mitosis, then associated with one SPB during anaphase B (Figure 4A). Similarly, *pmo25-35* cells showed Cdc7-YFP localized to the two SPBs at early mitosis, then concentrated to one SPB during anaphase B (Figure 4A). Thus, Pmo25 is not important for the SPB localization of Cdc7.

**Figure 4.**
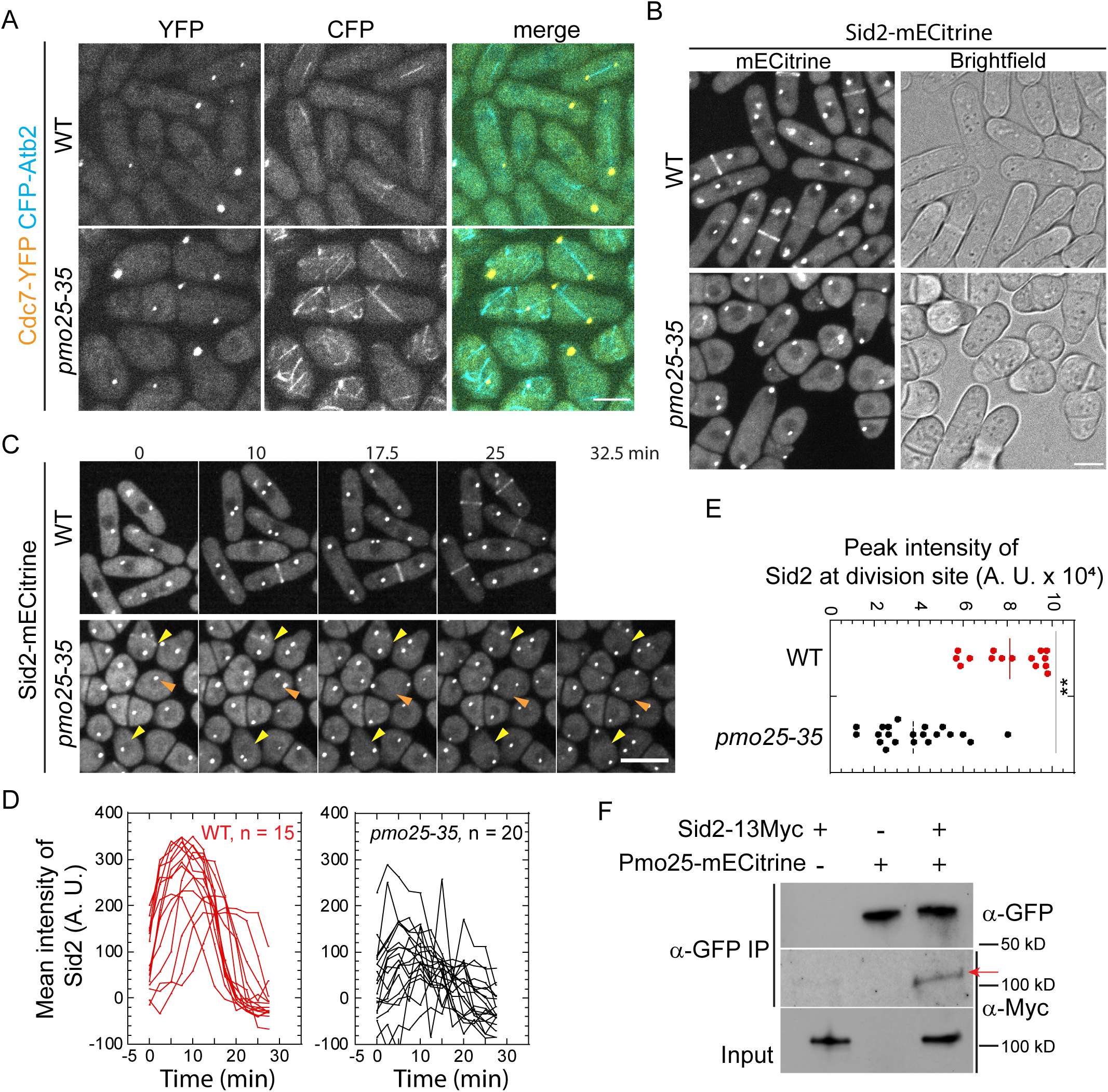
Pmo25 binds the SIN kinase Sid2 and is involved in Sid2’s recruitment to the division site. (A and B) Localization of the SIN kinases Cdc7 (A) and Sid2 (B) in *pmo25-35* mutant. Cells of WT (JW10070 and JW10139) and *pmo25-35* mutant (JW10063 and JW10138) expressing Cdc7-YFP or Sid2-mECitrine were grown at 25°C for ∼36 h, then shifted to 36°C for 4 to 5 h before imaging at 36°C. (C-E) Time course in minutes (C), mean intensity (D), and peak intensity (E) of Sid2-mECitrine at the division site. Cells (JW10138 and JW10139) were grown at 25°C for ∼36 h, then shifted to 36°C for 5 h before imaging at 36°C. In (C), yellow arrowheads: cells with weak Sid2 signal at the division site; orange arrowheads: almost no Sid2 signal at the division site during cytokinesis. (D) Time 0 is the start of the movie. (E) **p<0.0001. (F) Co-IP of Pmo25 with Sid2. Extracts of cells expressing Pmo25-mECitrine and/or Sid2-13Myc (YDM514, JW7943 and JW10082) were immunoprecipitated with anti-GFP polyclonal antibody, then detected by anti-Myc or anti-GFP monoclonal antibodies, respectively. The arrow marks the Sid2-13Myc band. Bars, 5 μm.

Surprisingly, the division-site localization of Sid2, the downstream kinase of the SIN pathway, was significantly affected in *pmo25* mutant cells (Figures 4B-4E, Movies 6 and 7). In WT cells, 5 h after shifting to 36°C, Sid2-mECitrine was still recruited to the division site during the late cytokinesis as reported (Figures 4B and 4C, Movie 6). In contrast, in *pmo25-35* cells at 36°C, the division-site recruitment of Sid2-mECitrine was significantly decreased (Figures 4B and 4C, Movie 7). We also found that some round *pmo25-35* cells, which may indicate a more severe Pmo25 activity defect, showed no detectable Sid2 signal at the division site (Figures 4B and 4C, Movie 7). To confirm the effect of Pmo25 on Sid2 distribution at the division site, we measured Sid2-mECitrine intensity at the division site in time-lapse movies. In WT cells, the Sid2 recruitment to the division site began before spindle break-down (indicated by Sid2 SPBs movement); and Sid2 rapidly reached peak intensity. However, in *pmo25-35* cells, the division-site recruitment of Sid2 was significantly compromised, with a ∼50% decrease in peak Sid2 intensity compared to WT cells (Figures 4D and 4E). Consistently, the Sid2 binding partner and regulatory subunit Mob1 showed similar defects in division site recruitment in *pmo25-35* cells (Figures S2A and S2B). The dependence of the Sid2 division-site localization on Pmo25 suggested they may interact physically with each other. Indeed, after immunoprecipitating Pmo25 with anti-GFP antibody, Sid2 was detected in cells co-expressing Pmo25-mECitrine and Sid2-13Myc (Figure 4F). Taken together, these results suggest that Pmo25 associates with Sid2 kinase and plays a role in the recruitment of the Sid2/Mob1 kinase complex to the division site during cytokinesis.

The role of Pmo25 in the recruitment and/or maintenance of Sid2 at the division site and their physical interactions in co-IP suggested that they must overlap temporally and spatially in *S. pombe* cells, which had not been examined before. Therefore, we tested their spatiotemporal relationship using confocal, time-lapse, and SoRa (Super Resolution by Optical Pixel Reassignment) high spatial resolution imaging in cells expressing both Sid2-mEGFP and Pmo25-tdTomato (Figures 5 and S3; and Movie 8). As reported before, Sid2 localized to one SPB during interphase and then to both SPBs and the division site during mitosis and cytokinesis^26,27,94-98^. Pmo25 localized to both SPBs transiently during early anaphase, then only to the new SPB with active Cdc7 kinase^33,34^. Sid2 and Pmo25 colocalized in the new SPB from mid to late anaphase until Pmo25 disappeared from the SPB near the end of contractile ring constriction in cells with a closed septum (Figures 5A and S3, and Movie 8). Sid2 was observed to form a discrete ring at the division site several minutes before Pmo25 during late anaphase (Figures 5A and S3; and Movie 8). Interestingly, Pmo25 appeared as a diffuse band on both sides of Sid2 at the division site first and then coalesced into a discrete ring that colocalized with Sid2 (Figures 5A and S3; and Movie 8). Sid2 and Pmo25 spread to the whole division plane during contractile-ring constriction and septum formation to form a washer-like structure and then a disc (Figure 5A). Sid2 faded away from the division site several minutes after septum closure and at least 10 minutes after Pmo25 disappeared from the new SPB (Figure S3). Pmo25 stayed at the division site until daughter-cell separation (Figure 5A). High-resolution SoRa microscopy confirmed that Pmo25 formed double discs^33,34^, which correspond to the two new layers of the plasma membrane at the division site (Figure 5B). These results were confirmed by using cells expressing both Sid2-mEGFP and Pmo25-mScarlet-I. Collectively, these data indicate that Sid2 and Pmo25 colocalize at the new SPB and the plasma membrane at the division site for >30 minutes, which supports their physical and functional interactions.

**Figure 5.**
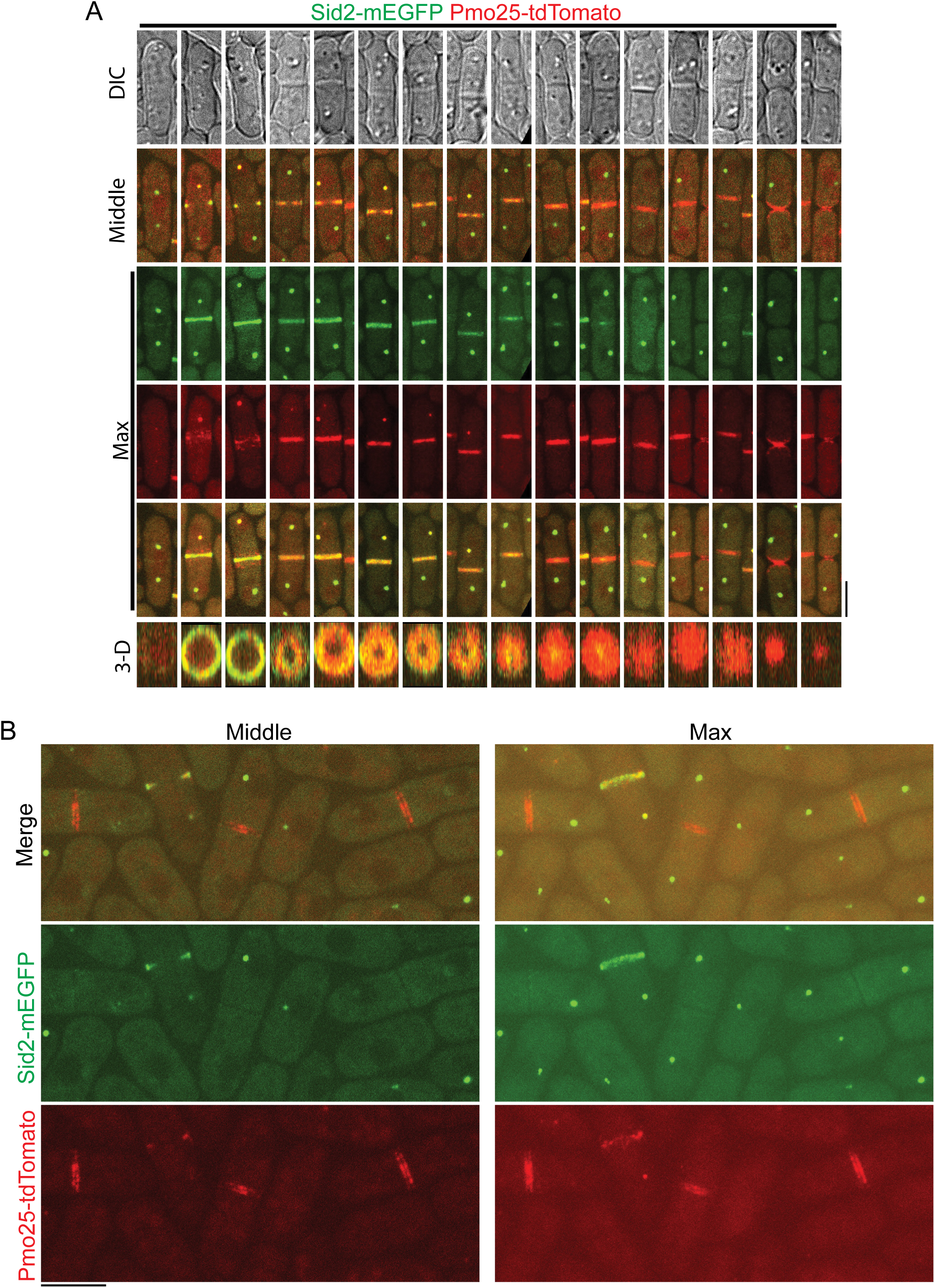
Colocalization of Sid2 and Pmo25 at the SPB and the division site during cytokinesis. (A) Colocalization of Sid2 and Pmo25 at the SPB and the division site. Single time-point images of cells expressing both Sid2-mEGFP and Pmo25-tdTomato (JW10302) are ordered chronologically based on septum morphology and Sid2/Pmo25 signals. DIC, middle focal plane, maximal intensity projection (19 slices with 0.3 μm spacing), and 3-D projections are shown. Cells were grown at 25°C for ∼48 h before imaging. (B) SoRa imaging of Pmo25 and Sid2 at the division site in the middle focal plane and max projections. Cells expressing Sid2-mEGFP and Pmo25-tdTomato (JW10302) were grown at 25°C for ∼48 h before imaging. Bars, 5 μm.

### Pmo25 is involved in the recruitment of the glucanase Eng1 to the division site during cytokinesis

Our data show that cytokinesis, septum formation, and SIN signaling are impaired in *pmo25* mutants (Figure 1); therefore, we examined whether *pmo25* mutants also affect glucan synthases and/or glucanases that are essential for successful cytokinesis and septation. First, we tested the localizations of the glucan synthases Bgs1, Bgs4, and Ags1, which assemble a three-layer septum composed of mainly α- and β-glucans during cytokinesis^67-69^. Rlc1-tdTomato was used as the contractile-ring marker. As expected^67-69^, GFP-Bgs1, GFP-Bgs4, and Ags1-GFP are transported by secretory vesicles and concentrated to the plasma membrane at the growing cell tips during interphase and recruited to the division site during the contractile ring maturation and constriction, and remain at the division site until daughter-cell separation (Figure 6A). In *pmo25-21* mutant cells, all three glucan synthases still localized to vesicles, cell tips, and the division site at comparable levels to WT cells at both 25°C and 36°C (Figure 6B). Thus, Pmo25 is not critical for recruitment of the glucan synthases.

**Figure 6.**
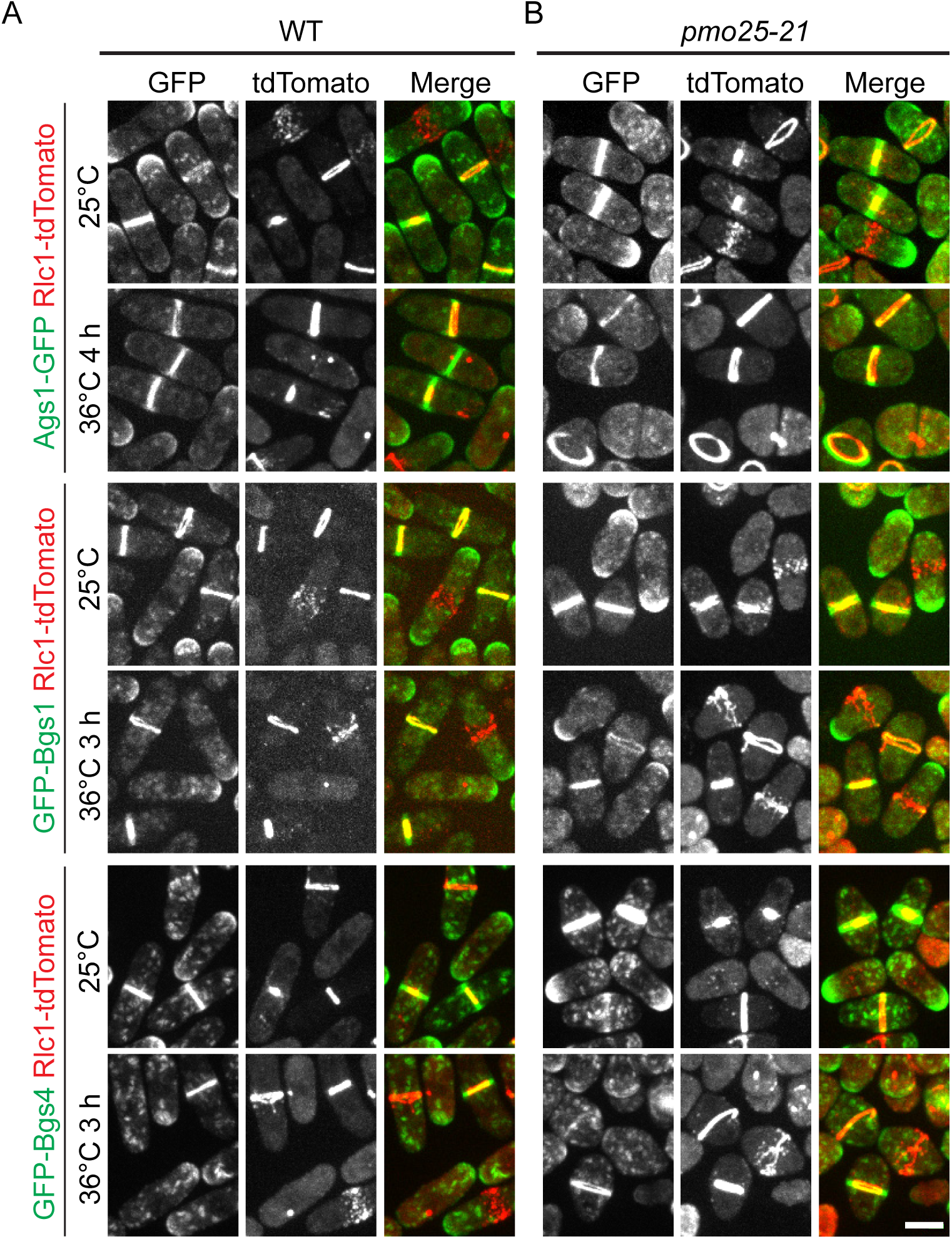
Localization of the glucan synthases Ags1, Bgs1, and Bgs4 in *pmo25-21* mutant. (A and B) Localization of Ags1, Bgs1, and Bgs4 in (A) WT and (B) *pmo25-21* mutant. Rlc1-tdTomato marks the contractile ring. Cells were grown at 25°C for ∼36 h then imaged, or shifted to 36°C for 3 h or 4 h before imaging at 36°C. Bar, 5 μm.

Next, we examined the distribution and intensity of the endo-1,3-β-glucanase Eng1 at the division site, which is involved in primary septum digestion to separate the two daughter cells after cytokinesis^76,77,99-101^. Eng1 is concentrated to the division site during late cytokinesis in WT cells as reported^76,77,99-101^ (Figures 7A and 7B). However, in *pmo25-21* cells, the division site level of Eng1 was significantly decreased even at 25°C. Because the division site signals of Eng1-mNeonGreen were weaker even in WT cells after shifting to 36°C compared to 25°C (Figures 7A and 7B), we measured Eng1-NeonGreen intensities during time lapse movies at 25°C (Figures 7C and 7D). In *pmo25-21* mutant cells, the peak levels of Eng1 at the division site were significantly lower than those in WT cells, and the maximal recruitment of Eng1-NeonGreen to the division site also took longer in *pmo25-21* cells (Figures 7C and 7D). Moreover, Eng1 stayed longer at the division site (Figure 7D), maybe due to the cytokinesis delay in *pmo25-21* cells. These data indicate that Pmo25 plays a role in the recruitment of glucanase Eng1 to the division site, which contributes to the delay of daughter-cell separation in *pmo25* mutants. Thus, Pmo25 not only plays a role in contractile-ring stability but also in daughter-cell separation.

**Figure 7.**
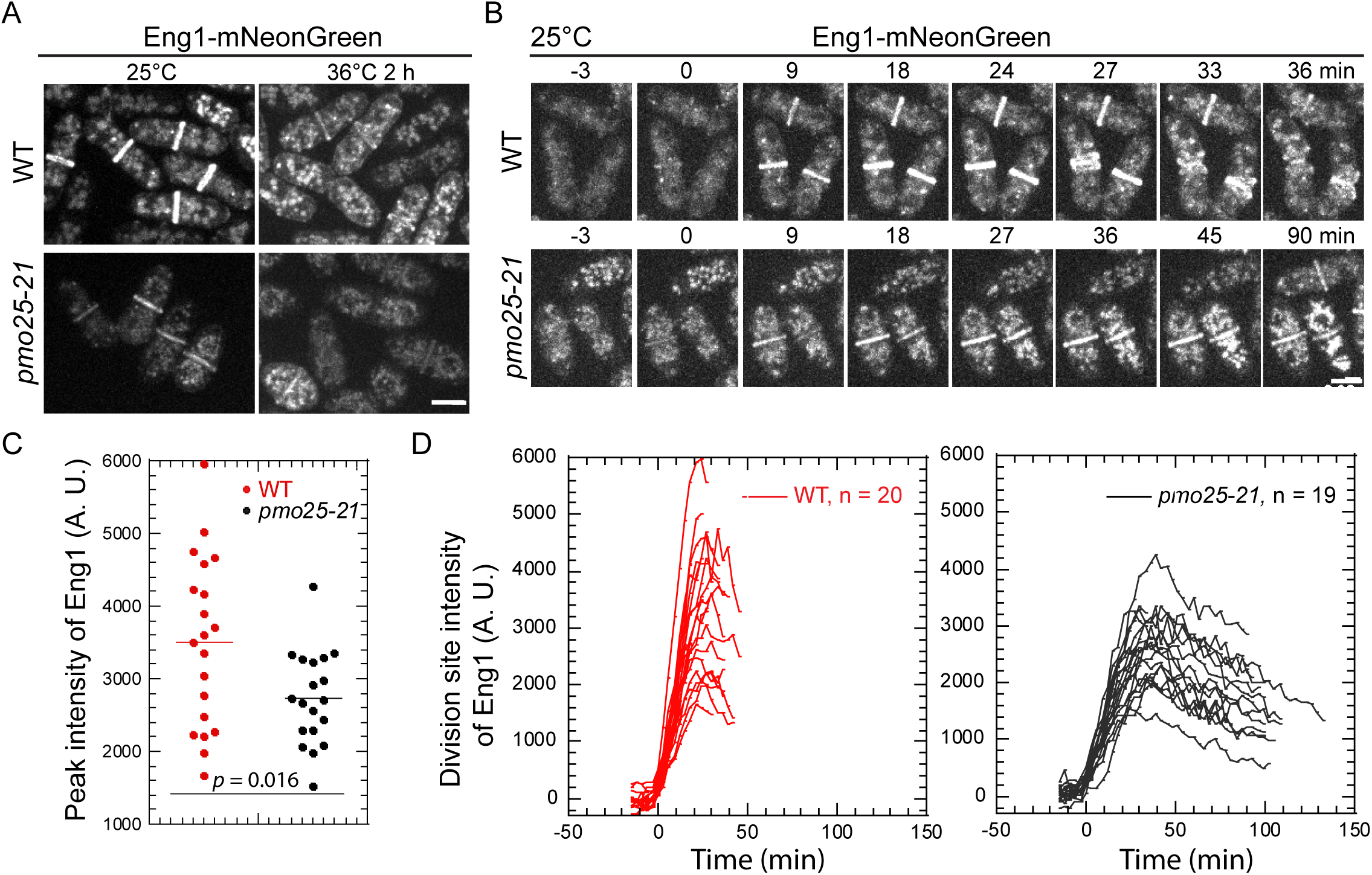
*pmo25-21* mutant affects β-glucanase Eng1’s recruitment to the division site. (A) Localization of Eng1 in WT (JW8489) and *pmo25-21* mutant (JW8488). Cells were grown at 25°C for ∼36 h then imaged, or shifted to 36°C for 2 h before imaging at ∼36°C. (B) Selected micrographs of Eng1 time course (in min) in WT (JW8489) and *pmo25-21* mutant (JW8488) grown at 25°C for ∼36 h before imaging. Time 0 marks Eng1 appearance at the division site. (C and D) Eng1 peak intensity (C) and time course of its mean intensity (D) at the division site in WT (JW8489) and *pmo25-21* cells (JW8488) grown at 25°C as in (B). (D) Time 0 marks Eng1 appearance at the division site and cells were followed until daughter-cell separation or the end of the movie. Bars, 5 μm.

### Pmo25 is a binding partner of Ync13

The Munc13/UNC-13 protein Ync13 in *S. pombe* plays an essential role in cell-wall integrity during cytokinesis, but its binding partners were previously unknown^58^. Our effort on identifying the binding partner of Ync13 converges on Pmo25. To identify binding partners of Ync13, we carried out yeast-two-hybrid screens. As full length Ync13 showed high levels of auto-activation in the *HIS3* and *URA3* reporter assays, Ync13 NH_2_-(1-590 aa) and COOH-terminus (591-1237 aa) were used as baits. A (19-329)-aa fragment of Pmo25 (SPAC1834.06c, full length: 329 aa) was found to exhibit strong positive interaction with Ync13 COOH-terminus (591-1237 aa), but not with its NH_2_-terminus (1-590 aa) (Figure 8A).

**Figure 8.**
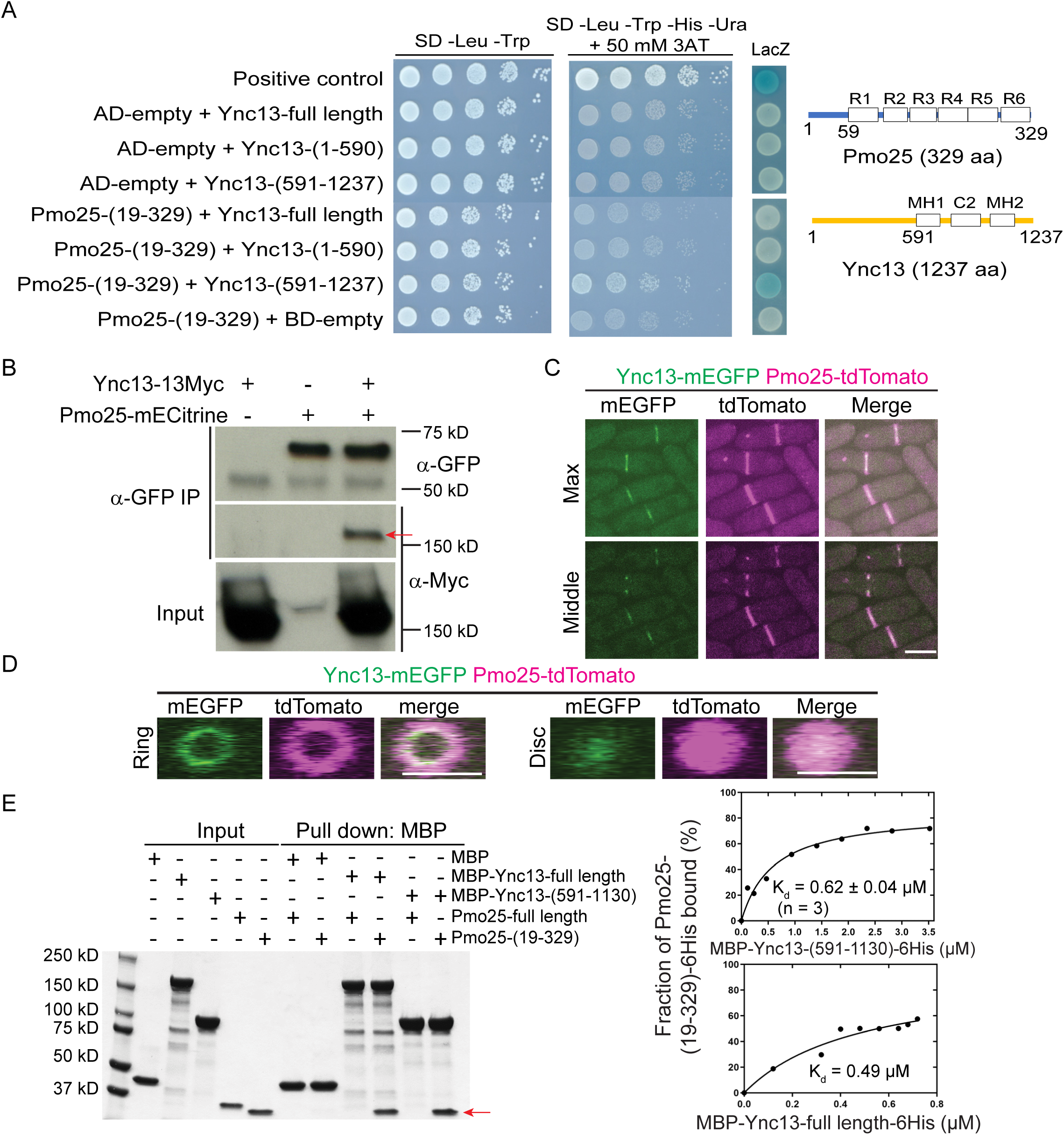
The Munc13/UNC-13 protein Ync13 interacts with the MO25 protein Pmo25. (A) Interaction between Ync13 and Pmo25 in yeast two hybrid assays. Left, different combinations of AD-Pmo25 and BD-Ync13 fragments were transformed into *S. cerevisiae* MaV203 cells. Interaction was detected by 10x serial dilution on quadruple dropout medium with 3-aminotriazole (3AT) and LacZ coloration. Right, Domain structures of Pmo25 and Ync13 ^33,45,58,128^. Pmo25 has six armadillo-like helical repeats (R1-R6). Ync13 has C2 domain and two Munc13 homology domains (MH). Numbers indicate the amino acids. (B) Co-IP of Ync13 and Pmo25. Cell extracts were immunoprecipitated with anti-GFP antibody. Precipitated and input samples were then detected using monoclonal antibodies against Myc and/or GFP. The arrow marks the Ync13-13Myc band. (C and D) Colocalization of Pmo25 and Ync13 in max projection and middle focal plane (C) and 3D projection (D). Cells (JW7970) expressing Pmo25-tdTomato and Ync13-mEGFP were grown at 25°C for ∼36 h before imaging with 0.4 μm spacing for 16 slices. Bars, 5 μm. (E) In vitro binding assay of Ync13 and Pmo25. Ync13 full length (1237 aa), Ync13 C terminal 591-1130 aa, Pmo25 full length (329 aa), and Pmo25 C terminal 19-329 aa were purified from *E. coli*. Binding assay was performed with MBP pull down. Left, protein samples analyzed by SDS-PAGE and stained with Coomassie Blue, the arrow marks the Pmo25-(19-329) band. Right, the curve fits and *K_d_* values for Pmo25-(19-329) with Ync13-(591-1130) and Ync13-full length, respectively.

To confirm the Pmo25-Ync13 interaction in *S. pombe* cells, we tagged Pmo25 and Ync13 at their COOH-terminus with mECitrine and 13Myc, respectively. After immunoprecipitating Pmo25 with anti-GFP antibody, Ync13-13Myc was pulled down from the cells co-expressing Pmo25-mECitrine and Ync13-13Myc, indicating that Pmo25 and Ync13 interact in fission yeast cells (Figure 8B). In addition, Ync13-mEGFP and Pmo25-tdTomato colocalized at the division site, though Ync13 was less abundant and more concentrated at the leading edge of the cleavage furrow (Figures 8C and 8D), which was confirmed using cells expressing Ync13-mECitrine and Pmo25-mScarlet. Unlike Pmo25, Ync13 did not localize to the SPBs (Figure 8C). To test whether Pmo25 interacts with Ync13 directly, we purified MBP-tagged full length Ync13 and Ync13 COOH-terminus (591-1130 aa), and His-tagged Pmo25 full length and Pmo25 COOH-terminus (19-329 aa) from *E. coli*. In vitro binding assays showed that neither Ync13 full length nor Ync13 C-terminus (591-1130 aa) interacted with Pmo25 full length, but both interacted with Pmo25-(19-329 aa), with *K_d_* values of 0.49 µM and 0.62 µM, respectively (Figure 8E). Together, these results indicate that Pmo25 is a direct binding partner of Ync13 in fission yeast.

The absence of residues 1-18 of Pmo25 promotes the interaction between Pmo25 and Ync13 in yeast two hybrid and in vitro binding assay. We constructed the *pmo25 (aa19-329)* mutant with residues 1-18 of *pmo25* deleted by gene targeting to test whether these residues are important for Pmo25 localization and function^116^. The mutant, expressed under its native promotor and at its endogenous chromosomal locus, had no obvious phenotypes in cell polarity or cytokinesis from 25 to 36°C (Figure S4A and data not shown). The localization of Pmo25(aa19-329) to the SPB and the division site also resembled the full length Pmo25 (Figure S4A). AlphaFold3^131^ was used to predict the structure of full length Pmo25 (pTM = 0.91) and Pmo25(aa19-329) (pTM = 0.92) with high confidence in most domains except the unstructured Pmo25(aa1-18). These two structures showed a high degree of similarity with an RMSD (Root Mean Square Deviation) of only 0.156 Å across the 311 aa when superimposed. Thus, Pmo25(aa1-18) is likely unstructured, and its deletion has no obvious effects on the overall Pmo25 structure (Figure S4B). Future studies are needed to elucidate the functional and regulatory significance of Pmo25(aa1-18).

### Pmo25 affects Ync13 accumulation at the division site

Apart from its role in the SIN pathway, Ync13 also affects septum integrity by regulating the distribution of glucan synthases and glucanases at the division site^58^. To determine the functional relationship between Pmo25 and Ync13, we tested their localization dependency. Ync13-mECitrine localized to the cell tips during interphase and the division site during mitosis and cytokinesis; and Pmo25-mECitrine localized to the SPBs and the division site during mitosis and cytokinesis (Figures 9A-9D), consistent with Pmo25-tdTomato and the previous reports^33,34,58^. When the SPB marker Sad1-mCherry was used to monitor mitotic progress, >67% of cells with two SPBs showed Ync13 localization at the division site, and the mean SPB distance was ∼5 μm. In contrast, <50% of cells with two SPBs showed Pmo25 localization at the division site, and the mean SPB distance was ∼8 μm in cells with Pmo25 at the division site, indicating that Ync13 appears at the division site earlier than Pmo25 (Figure 9A). However, *ync13Δ* cells had no obvious defect in Pmo25 localization at the division site (Figure 9B and 9C). Deletion of *ync13* leads to cell lysis in YE5S (<u>Y</u>east <u>E</u>xtract with <u>5</u> <u>S</u>upplements) rich medium without the osmotic stabilizer sorbitol^58^. Pmo25 localized to the SPBs and the division site at comparable levels in *ync13Δ* cells and WT cells (Figure 9B and 9C). However, although Ync13 still localizes to the cell tips and the division site in *pmo25-2* cells at the restrictive temperature of 36°C, its mean intensity at the division site increased dramatically (Figure 9D and 9E). Collectively, Pmo25 directly interacts and partially colocalizes with Ync13 and is important for the Ync13 level at the division site.

**Figure 9.**
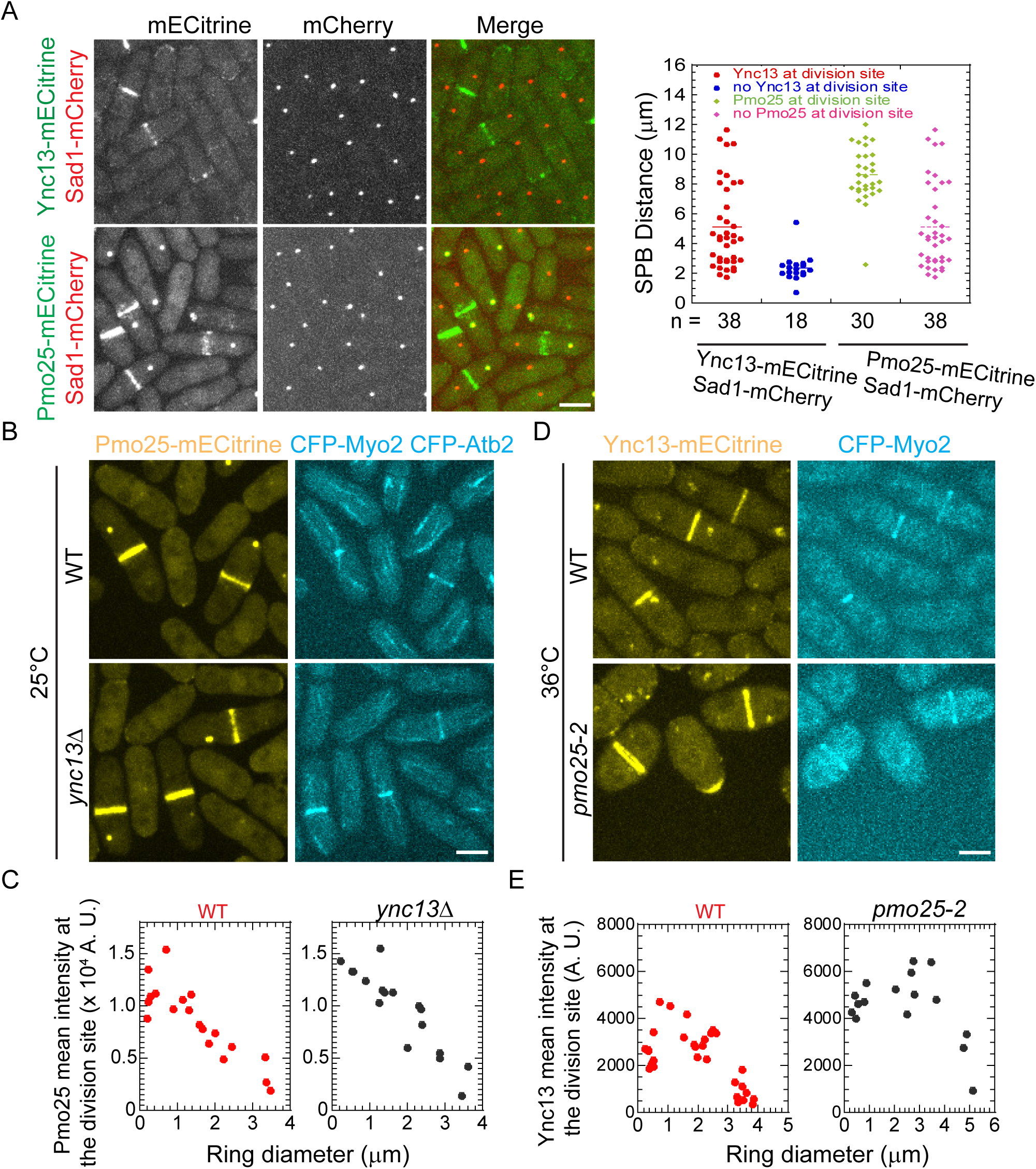
Timing of Pmo25 localization to the division site and its effect on the Ync13 level during cell division. (A) Micrographs (left) and quantifications (right) of Pmo25 (JW7968) and Ync13 (JW5814) localizations in the cells with Sad1-mCherry as a cell-cycle marker. Right, the timing of Pmo25 and Ync13 appearance at the division site relative to the distance between two SPBs. (B and C) Localization and division site intensity of Pmo25 in WT and *ync13Δ* mutant. Cells of WT (JW8174) and *ync13Δ* mutant (JW8396) were grown in YE5S + 1.2 M sorbitol at 25°C for ∼36 h, then washed into YE5S and grown for 3 h before imaging. The division site intensity of Pmo25 was measured in SUM projection and plotted versus Myo2 ring diameter. (D and E) Localization and division site intensity Ync13 in WT and *pmo25-2* mutant. Cells of WT (JW8135) and *pmo25-2* mutant (JW8136) were grown at 25°C for ∼36 h, then shifted to 36°C for 2 h before imaging. The mean intensity of Ync13 in the ROI (Region of Interest) at the division site was measured in SUM projection and plotted versus Myo2 ring diameter.

## DISCUSSION

In this study, we revealed that Pmo25, an MO25 protein, plays important roles in contractile-ring stability, SIN signaling, and daughter-cell separation during cytokinesis. We found that Pmo25 directly interacts with the Munc13/UNC-13 protein Ync13 and is also a candidate binding partner of the myosin-II light chain Cdc4 and the NDR kinase Sid2. Thus, this indicates that Pmo25 may connect different stages of cytokinesis in addition to its known roles in cell polarity. Because all these proteins are highly conserved during evolution^4,37,38,42,43,45,46,52,54,58,65^, our data provide insight into their potential interactions and functions in cell morphogenesis and cell division in other systems.

### Contractile-ring stability and the SIN signaling are disrupted in *pmo25* mutants during cytokinesis

Previous studies found that Pmo25 is a component of the MOR network and is essential for activation of the Orb6 kinase during cell polarization and morphogenesis^33,34,53^. Pmo25 mainly localizes to the SPB with active SIN signaling during mitosis and to the division site during cytokinesis^33,34,53^. It mediates the signaling connection between the MOR and SIN pathways^33,34,53^. It was reported that *pmo25* mutants are defective in cell separation after septum formation with an increased septation index and some multiseptated cells^33,34^. However, the molecular mechanism of how Pmo25 regulates cytokinesis remained a mystery. In budding yeast, the MO25 protein Hym1 has been shown to be involved in cell separation but not in contractile-ring stability^4,47,102^. Here we presented several lines of evidence that Pmo25 is important for the stability and maintenance of the contractile ring. First, in *pmo25* temperature-sensitive or deletion mutants and in combination with the *bgs1/cps1-191* mutant, the contractile ring frequently collapses, frays, deforms, or disintegrates before or during constriction. Second, Pmo25 physically interacts with Cdc4 in yeast two hybrid and Co-IP assays and most mYFP-Cdc4 rings collapse during cytokinesis. Cdc4 is an essential protein in contractile-ring stability and maintenance as a binding partner of the myosin-IIs Myo2 and Myp2, the myosin-V Myo51, and the IQGAP Rng2^82,103-107^. Third, *pmo25* mutants have strong genetic interactions with all the tested mutations affecting contractile-ring stability and maintenance such as mutations in *cdc12*, *rng2*, *rlc1, myo2* and *cdc4,* even at permissive temperatures for *pmo25* mutants (Table 1). Their double mutants display the typical phenotypes of classic cytokinesis mutants, such as incomplete and disorganized septa and multiple nuclei while maintaining the rod shape (Figure S1). Although *pmo25* temperature-sensitive or deletion mutants have severe polarity defects similar to the *orb6* mutant, the contractile ring is less stable and more defective in *pmo25* than in the *orb6* mutant (Movie 1). Thus, we conclude that Pmo25 plays an important role in contractile-ring stability before and during its constriction.

How does Pmo25 affect the contractile-ring stability? One possibility is that Pmo25 interacts with Cdc4 to stabilize the various complexes of Cdc4 with other ring proteins, which is supported by the strong genetic interactions between mutants of *pmo25* and the binding partners of Cdc4 such as *myo2* and *rng2* (Table 1). The other possibility is that Pmo25 affects ring stability through the SIN pathway, which is known to regulate contractile-ring stability and maintenance^8,12,24,89^. The NDR kinase Sid2 is the most downstream kinase in the SIN pathway. In this study, we found that the levels of Sid2 and its regulatory subunit Mob1 at the division site are dramatically reduced in *pmo25* mutants. Their peak intensity and kinetics of recruitment at the division site are both severely compromised. In addition, Sid2 and Pmo25 colocalize at the division site and the SPBs during cytokinesis, and they interact in Co-IP assays, which had not been tested previously. Thus, it is possible that Pmo25 is one of the players that regulate the contractile ring through the SIN pathway during cytokinesis. However, it is also possible that Pmo25 functions in the *bgs1/cps1-191* contractile-ring checkpoint rather than in ring stability per se^132,133^. Future experiments are needed to distinguish these possibilities. It would be interesting to test if MO25 interacts with the tumor suppressor NDR kinase STK38, the Sid2 homolog in mammalian cells^108-112^. MO25 forms a heterotrimeric complex with the pseudokinase STRAD and the tumor suppressor liver kinase 1 (LKB1). The LKB1-STRAD-MO25 complex stabilizes a closed conformation of STRAD and triggers LKB1 nucleocytoplasmic shuttling and activation^36-39^. It seems that Pmo25 also forms a heterotrimeric complex with the Sid2-Mob1 kinase complex during cytokinesis. Thus, it would be interesting to test if MO25 proteins interact with the NDR kinase in the Hippo pathways to regulate cytokinesis in animal cells.

### Pmo25 interacts with Ync13 to modulate the recruitment of the glucanase Eng1 at the division site

We showed that Pmo25 and Ync13 physically interact with each other by yeast two-hybrid, Co-IP, and in vitro binding assays. Their interaction is direct with a relatively strong affinity (K_d_ < 1 µM). The Pmo25 level at the division site is not affected by *ync13Δ*. By contrast, Ync13 fluorescence intensity at the division site increases by almost 50% in *pmo25* mutants. Further studies are needed to elucidate the mechanism by which the Ync13 level at the division site increases in *pmo25* mutant cells. We investigated the functions of the Ync13-Pmo25 interaction in fission yeast which demonstrated that Ync13 and Pmo25 partially colocalize on the plasma membrane at the division site with Ync13 being more concentrated at the leading edge. Because *ync13* mutants have no effects on contractile-ring stability or maintenance and the Pmo25 level at the division site is normal without Ync13, the Ync13-Pmo25 interaction is less likely to be involved in regulating the contractile ring. It is more likely this interaction is important for recruiting the glucanase Eng1 for daughter-cell separation via exocytosis. Consistently, the Eng1 level and recruitment at the division site are compromised in both *pmo25* (Figure 7) and *ync13* mutants ^58^. Thus, both Ync13 and Pmo25 play important roles in septum formation and daughter-cell separation. However, although both *ync13* and *pmo25* mutants are defective in cytokinesis and undergo cell lysis, their morphologies are not identical. Unlike Ync13, Pmo25 localizes to the SPBs and *pmo25* mutant cells lose polarity, while *ync13* mutant cells do not. Thus, Ync13 and Pmo25 must also have independent functions besides working together, which need to be examined in future studies.

Our data suggests several novel functions of Pmo25 that modulate multiple cellular processes such as contractile-ring stability and constriction, exocytic trafficking, and cell separation. We predict that Pmo25 may exist in several protein complexes to recruit its binding partners to the cell-division site and SPBs at different times. It is also possible that Pmo25 acts as a scaffold to link these sub-complexes together into a mega-complex, which may coordinate cytokinesis and the cell cycle more efficiently. This is supported by the fact that Pmo25 and its binding partners colocalize at the division site in spatiotemporally overlapping and distinct patterns. How these modules coordinate various cellular events and whether Pmo25 regulates them in different ways, requires further investigation.

### Limitations of the study

In this study, we propose that Pmo25 functions as a coordinator with several partners or complexes during cytokinesis. However, the detailed molecular mechanisms were not explored. Whether Pmo25 interacts with Cdc4 and Sid2 directly or indirectly has also not been tested by in vitro binding assays. Their in vivo localizations at the division site show spatiotemporal overlapping, though it is unknown how Pmo25 is involved in different complexes. More studies are needed to determine if Pmo25 regulates the contractile-ring stability through proteins other than Cdc4 and Sid2. Pmo25 affects Sid2-Mob1 complex recruitment to the division site, and this might be independent from the MOR pathway^31,32,53^. In future studies, structure and function analyses are needed to dissect what domains and residues contribute to the interactions between Pmo25 and different proteins as well as the physiological roles of these interactions. The dynamics of Ync13 in *pmo25* mutants needs to be tested using FRAP assays, which is challenging because Ync13 signal is weak and Ync13 is highly dynamic at the division site with half times of 2 to 3 seconds^58^. Pmo25 and Ync13 interact with each other and partially colocalize at the division site. It is unknown why Pmo25 (19-329) but not full length Pmo25 interacts with Ync13. The first 18 residues of Pmo25 are upstream of the armadillo-like helical repeats and have no obvious motif. We hypothesize that residues 1-18 (maybe together with the adjacent regions) of Pmo25 may autoinhibit the binding domain of Pmo25 with Ync13. This possible autoinhibition could be regulated in the fission yeast cells because the full length Pmo25 binds to Ync13 in co-IP from *S. pombe* cell extract. Future studies are needed to investigate the function of the NH2 terminal part of Pmo25 and those of the armadillo-like repeats in such potential autoinhibition in Pmo25.

## RESOURCE AVAILABILITY

### Lead contact

Requests for further information, data, and reagents should be directed to and will be fulfilled by the lead contact, Jian-Qiu Wu

### Materials availability

Yeast strains and plasmids generated in this study will be available upon request.

### Data and code availability

This paper does not report original code. Any additional information required to reanalyze the data reported in this paper is available from the lead contact upon request.

## ACKNOWLEDGMENTS

We thank Mohan Balasubramanian, Damian Brunner, Juan Carlos, Fred Chang, Noah DiFilippo, Kathy Gould, Dai Hirata, Kazunori Kume, Dan McCollum, Tom Pollard, Beatriz Santos, Ken Sawin, and Yihua Zhu for yeast strains and/or plasmids; Davinder Singh, Clara Sablak, Nick Ricottilli, and Lexi Waligura for technical support; Anita Hopper for equipment; and current and former members of the Wu lab for helpful discussions. S.Z. was supported by Pelotonia Postdoctoral Fellowship Program and E.G.G. by Pelotonia Undergraduate Fellowship Program from The Ohio State University. J.R.G. was supported by the Cellular, Molecular, and Biochemical Sciences Training Program (T32 GM141955) from the National Institutes of Health. The work was supported by the National Institute of General Medical Sciences of the National Institutes of Health (grants R01 GM118746 and GM118746-06S1 to J.-Q.W.). The content is solely the responsibility of the authors and does not necessarily represent the official views of Pelotonia Fellowship Program or The Ohio State University or the National Institutes of Health.

## AUTHOR CONTRIBUTIONS

Conceptualization, Y.Y. and J.-Q.W.; Methodology, Y.Y., S.Z., and J.-Q.W.; Investigation, Y.Y., S.Z., J.R.G., A.H.O., E.G.G., and J.-Q.W.; Formal Analysis, Y.Y., S.Z., J.R.G., A.H.O., E.G.G.; Validation, Y.Y., S.Z., A.H.O., J.R.G.; Visualization, Y.Y., J.R.G.; Writing – Original Draft, Y.Y., J.R.G., and J.-Q.W.; Writing –Review & Editing, Y.Y., J.R.G., E.G.G., and J.-Q.W.; Funding Acquisition, S.Z., E.G.G., J.R.G., and J.-Q.W.; Resources, J.-Q.W.; Supervision, J.- Q.W.

## DECLARATION OF INTRESTS

The authors declare no competing interests.

## STAR*METHODS

Detailed methods are provided in the online version of this paper and include the following:

- KEY RESOURCE TABLE
- EXPERIMENTAL MODEL AND SUBJECT DETAILS
- METHODS DETAILS

- Strains, genetics, and cellular methods
- Yeast two hybrid screen and assays
- IP and Western blotting
- Protein purification and in vitro binding assays
- Microscopy and data analyses
- QUANTIFICATION AND STATISTICAL ANALYSIS

## SUPPLEMENTAL INFORMATION TITLES AND LEGENDS

Supplemental information can be found with this article online at ???

Figures S1-S4, Table S1, and Movies S1 to S8.

## STAR*METHODS

### KEY RESOURCE TABLE

| REAGENT or RESOURCE | SOURCE | IDENTIFIER |
| --- | --- | --- |
| **Antibodies** |  |  |
| Rabbit anti-GFP | Novus Biologicals | NB600-308 |
| Anti-GFP/YFP monoclonal | Roche | 11814460001 |
| Anti-Myc | Santa Cruz Biotechnology | 9E10 |
| Anti-mouse IgG | Sigma | A4416 |
| **Chemicals and reagents** |  |  |
| Dynabeads | Invitrogen | 10004D |
| BL21(DE3)pLysS | Novagen | 694513 |
| Protease inhibitor cocktail cOmplete | Roche | 11873580001 |
| Amylose beads | New England Biolabs | 8021S |
| n-propyl-gallate | Thermo Fisher Scientific | P3130-100G |
| X-Gal | Thermo Fisher Scientific | BP1615-1 |
| Gelatin | Sigma-Aldrich | G2500-500G |
| BioMax MR film | Kodak | Z350370-50EA |
| pACT2 yeast two hybrid library | Lab stock | N/A |
| Talon metal affinity resin | Clontech | 635501 |
| SuperSignal west maximum sensitivity substrate | Thermo Fisher Scientific | 34096 |
| **Recombinant DNA** |  |  |
| pFA6a-mEGFP-kanMX6 | Lab stock | JQW85 |
| pFA6a-mYFP-kanMX6 | Lab stock | JQW86 |
| pFA6a-tdTomato-kanMX6 | Ken Sawin<sup>113</sup> | pKS392 |
| pFA6a-tdTomato-natMX6 | Ken Sawin<sup>113</sup> | pKS393 |
| pFA6a-mECitrine-kanMX6 | Lab stock | JQW228 |
| pFA6a-mNeonGreen-kanMX6 | Lab stock | JQW946 |
| pFA6a-mScarlet-I-kanMX6 | Lab stock | JQW1020 |
| **Software and algorithms** |  |  |
| Fiji Software | https://fiji.sc<sup>114</sup> | RRID:SCR_002285 |
| Volocity | PerkinElmer | N/A |
| NIS Elements | Nikon | N/A |

### EXPERIMENTAL MODEL AND SUBJECT DETAILS

#### Fission Yeast

Yeast strains used in this study are listed in Supplemental Table S1. Cells were woken up from -80°C stocks and grown on YE5S plates at 25°C for ∼2 d, and then fresh cells were inoculated into YE5S liquid medium and grown at 25°C for ∼36 h at log phase (diluted twice daily) before imaging except where noted. Detailed growth conditions for individual experiments are in figure legends.

##### Escherichia coli

The plasmids were transformed into BL21 (DE3) pLysS cells (Novagen, 694513) for protein expression. MBP-Ync13-full length-6His and MBP-Ync13-(591-1130)-6His were induced with 0.2 mM isopropyl-β-D-thiogalactoside (IPTG) at 17°C for 36–48 h. Pmo25-full-length-6His and Pmo25-(19–329)-6His were expressed with 0.5 mM IPTG at 25°C for 15 h.

### METHOD DETAILS

#### Strains, genetics, and cellular methods

Strains used in this study are listed in Supplemental Table S1. PCR-based gene targeting and genetic crosses were performed using standard methods^115,116^. All tagged genes are expressed under endogenous promoters and integrated into the native chromosomal loci except tagged *psy1*, *bgs1*, *bgs4*, *ags1*, which are at the *leu1* locus. Epitope-tagged Pmo25 proteins were functional as the cells expressing them resembled WT in both growth and morphology from 25 to 36°C. The *pmo25* deletion mutant was constructed in the diploid strain *h^+^/h^-^ rlc1-tdTomato-natMX6/rlc1-tdTomato-natMX6 leu1^+^::GFP-psy1/ leu1^+^::GFP-psy1 Patb2-mRFP-atb2/Patb2-mRFP-atb2 ade6-M210/ade6-M216 ura4/ura4* using the *hphMX6* marker. Cells in Supplemental Figure S1 were stained with 1/33 volume of 1 mg/ml Hoechst 33258 (bisbenzimide) for 10 min before imaging.

#### Yeast two hybrid screen and assays

Yeast two hybrid screen was carried out as previously described^117^. The fragments corresponding to Ync13 full length (1237 aa), NH_2_-terminal (1-590 aa), and COOH-terminal (591-1237 aa) were amplified from a cDNA library and cloned into BD (DNA binding domain) pGBT9 vector, and co-expressed in *S. cerevisiae* MaV203 strain with prey library pACT2 (lab stock) by sequential transformation. The selection was performed with reporter genes: *URA3^+^*, *HIS3^+^*, and *LacZ*. For X-gal overlay assay, fresh colonies from leucine and tryptophan selection plates were re-streaked on YPD (yeast extract-peptone-dextrose) plates and grew overnight. ∼8 ml chloroform per plate was used to permeabilize cells for 10 min and then dried for additional 2 min before overlay. The overlay solution was prepared in 25 ml PBS (pH 7.5) with 0.5% agarose and 500 µl X-gal (20 mg/ml stock in DMSO). The overlaid plates were incubated at 30°C and checked for the development of blue color every 30 min. Plasmid DNAs from the positive colonies were isolated and sequenced.

To test the interactions between Pmo25 and the contractile ring components, the full length Pmo25 cDNA was cloned into VP16 transcription activation domain (AD) vector, pVP16, and co-transformed into *S. cerevisiae* MaV203 strain with pGBT9 vector expressing full length Rng2, Cdc15, Cdc12, Myo2, and Cdc4^82,118,119^, respectively. The selection was performed with reporter genes as described above. All constructed plasmids were confirmed by Sanger sequencing.

#### IP and Western blotting

Co-IP and Western blotting were performed as previously described^82,120,121^. Yeast cells expressing tagged proteins under the control of native promoters were harvested at exponential phase, washed twice with ddH_2_O, frozen immediately with liquid nitrogen, and then lyophilized. Lyophilized cells were further ground into a homogeneous powder in mortar with liquid nitrogen. ∼100 mg cell powder for each sample was dissolved in IP buffer (50 mM 4-(2-hydroxyethyl)-1-piperazineethanesulfonic acid [HEPES], pH 7.5, 150 mM NaCl, 1 mM EDTA, 0.5% NP-40 [for Ync13 and Pmo25 IP, 0.5% Triton and 0.5% CHAPS were used], 0.1 mM Na_3_VO_4_, 1 mM PMSF, and EDTA-free protease inhibitor cocktail [cOmplete, Roche]) on ice, then centrifuged at 10,000 g for 20 min at 4°C. 30 μl of protein G covalently coupled magnetic Dynabeads (100.04D; Invitrogen, Carlsbad, CA) per sample was washed three times with cold phosphate-buffered saline (PBS) buffer (2.7 mM KCl, 137 mM NaCl, 10 mM Na_2_HPO4, and 2 mM KH_2_PO4, pH 7.4), and 5 µl of rabbit anti-GFP antibody (NB600-308; Novus Biologicals, Littleton, CO) per sample in PBS buffer was added to beads. After incubation for 1 h at room temperature, the beads were washed three times with PBS buffer and twice with the IP buffer. 300 µl cell extracts from centrifuged supernatants were mixed with antibody-coupled Dynabeads and incubated for 1.5 h at 4°C. The precipitated beads were washed twice with wash buffer I (50 mM HEPES, pH 7.5, 150 mM NaCl, 1 mM EDTA, 0.1% NP-40) and three times with wash buffer II (50 mM HEPES, pH 7.5, 150 mM NaCl, 1 mM EDTA), then dissolved in sample buffer and boiled for 5 min. After separation of proteins in SDS–PAGE, proteins were detected by Western blotting using monoclonal anti–GFP/YFP antibody (1:2500 dilution; Roche) or monoclonal anti-Myc antibody (9E10, 1:1000 dilution; Santa Cruz Biotechnology, Santa Cruz, CA).

#### Protein purification and in vitro binding assays

The MBP-TEV-GGSGGS fragment was first inserted into the pET21a vector upstream of the *Bam*HI site using Gibson assembly^122^ to generate the pET21a-MBP construct. The full-length Ync13 cDNA was subsequently cloned into the pET21a-MBP vector between the GGSGGS linker and the C-terminal 6×His tag via Gibson assembly. Similarly, full-length Pmo25 and its truncated form (residues 19–329) were cloned into the pET21a vector between the N-terminal T7 tag and the C-terminal 6×His tag using the same method. All constructs were verified by DNA sequencing.

MBP-Ync13-full length-6His, MBP-Ync13-(591-1130)-6His, Pmo25-full length-6His, and Pmo25-(19-329)-6His were purified with Talon metal affinity resin (635501; Clontech, Mountain View, CA) using extraction buffer (50 mM sodium phosphate, pH 8.0, 450 mM NaCl, 10 mM β-mercaptoethanol, 1 mM PMSF, and 10 mM imidazole) with EDTA-free protease inhibitor tablet (Roche) as described before^123^. Proteins were eluted with elution buffer (50 mM sodium phosphate, pH 8.0, 450 mM NaCl, 10 mM β-mercaptoethanol, 1 mM PMSF, and 200 mM imidazole). The purified proteins were then dialyzed into the binding buffer (137 mM NaCl, 2 mM KCl, 10 mM Na_2_HPO_4_, 2 mM KH_2_PO_4_, 0.5 mM dithiothreitol, and 10% glycerol, pH 7.4).

For in vitro binding assays between Ync13 and Pmo25, we incubated MBP-Ync13-full length-6His, MBP-Ync13-(591-1130)-6His or MBP-6His control with 500 μl amylose beads for 1 h at 4°C and then washed the beads eight times with 1 ml of the binding buffer each time to remove unbound proteins. Then Pmo25-full length-6His or Pmo25-(19-329)-6His was incubated with the 100 μl beads with bound MBP-Ync13 or MBP proteins for 1 h at 4°C. After 4 washes with 1 ml of the binding buffer each time, the beads were boiled with sample buffer for 5 min. Then the samples were run on SDS–PAGE gel and detected with Coomassie Blue staining.

To measure the Kd between Ync13-(full length/591-1130) and Pmo25-(19–329), we followed the described methods and guidelines^123,124^. Various concentrations of MBP-Ync13- (full length/591-1130)-6His immobilized on amylose beads were incubated with a fixed low concentration of Pmo25-(19–329)-6His for 1h at 4°C. After incubation, the beads were spun down at 1,000 g for 1 min, supernatant samples removed from the reactions were boiled with sample buffer, run on an SDS-PAGE gel, and detected with Coomassie Blue staining. Total amount of bead bound Pmo25-(19-329)-6His was calculated by subtracting the remaining proteins in the supernatant from the total input. The Kd was calculated using one site binding Equation in Graphpad Prism 9.5.0.

#### Microscopy and data analyses

For microscopy imaging, cells were woken up from -80°C stocks and grown on YE5S plates at 25°C for ∼2 d, and then fresh cells were inoculated into YE5S liquid medium and grown at 25°C for ∼36 h at log phase (diluted twice daily) before imaging except where noted. For cold-sensitive strains such as *rlc1Δ*, cells were woken up and grown at 32°C before shifting to lower temperatures before imaging. Microscopy sample preparations were carried out as described^80,87,120,125^. Briefly, cultured cells were collected by centrifugation at 3,000 to 3,500 rpm, washed once with EMM5S, and then washed with EMM5S containing 5 µM *n*-propyl-gallate to reduce autofluorescence and protect cells from free radicals during microscopy. Cells were spotted onto a slide with EMM5S containing 5 µM *n*-propyl-gallate + 20% gelatin, sealed with VALAP. For imaging *pmo25Δ* cells using confocal fluorescence microscope, dissected tetrads were grown on YE5S agar plate for ∼16 h, then a fraction of WT cells and all *pmo25Δ* cells (predicted by phenotype) were collected to separate positions by using glass needle on the tetrad dissection microscope, grew further on YE5S agar plate at 25°C. The piece of agar plate with the collected cells was cut out and placed face down on a 35-mm dish with a glass coverslip bottom, an 18 x 18 mm coverslip was covered on the top as well to slow down the drying of agar media and imaged directly. Hygromycin sensitivity was checked with the leftover WT cells.

All fluorescence microscopy was carried out at ∼23°C (except where noted) on a PerkinElmer spinning disk confocal system (UltraVIEW Vox CSUX1 system; PerkinElmer, Waltham, MA) on a Nikon Ti-E microscope with Hamamatsu EMCCD camera C9100-23B and Plan-Apo 100x/1.45 NA objective; or on a Nikon CSU-W1 SoRa spinning disk confocal microscope with Hamamatsu ORCA Quest qCMOS camera C15550 on Nikon Eclipse Ti2 microscope and Apo TIRF 100x/1.49 NA oil, Plan Apo λD 100x/1.45 NA oil, or Plan Flour 100x/1.30 NA oil objectives.

To observe *pmo25*Δ phenotype under DIC in Figure 2A, the YE5S agar plate with dissected tetrads was cut out and placed on a glass slide, after covered with a glass coverslip, sample was sealed with VALAP and imaged with a 100×/1.4 numerical aperture (NA) Plan-Apo objective lens on a Nikon Eclipse Ti inverted microscope (Nikon, Melville, NY) equipped with a Nikon cooled digital camera DS-QI1. Cells for Figure S1 were also imaged using the same microscope on a slide with EMM5S containing 5 µM *n*-propyl-gallate + 20% gelatin.

Images and data were collected and analyzed by Volocity, NIS Elements, and Fiji software. The fluorescence intensity at the division site was measured in the images that were projected with sum intensity of 0.5 µm-spaced Z-slices as described previously^126,127^. ROI (region of interest) covering the signal at the division site was drawn to measure the mean intensity, and ∼2× ROI was used to measure and calculate background intensity^58,118,120^. Micrographs shown in the figures are maximum projections except where noted. Statistical tests were performed using a two-tailed Student’s *t* test.

#### AlphaFold3 modeling for structural predictions

Utilizing AlphaFold3, the deep-learning protein structural prediction tool, we predicted the structures of full length Pmo25 and Pmo25 (aa19-329)^131^. The amino acid sequence for Pmo25 was downloaded from Pombase (https://www.pombase.org/gene/SPAC1834.06c) <u>and inputted into AlphaFold3</u> (https://alphafoldserver.com). For Pmo25(aa19-329), the first 18 amino acids were deleted. All sequences were submitted in triplicate and had the same structures and predicted template modeling (pTM) values, which is an assessment of the accuracy of the generated structure^134,135^. The results were downloaded and analyzed via Chimaera-UCSF^136^.

---

**Figure S1.**
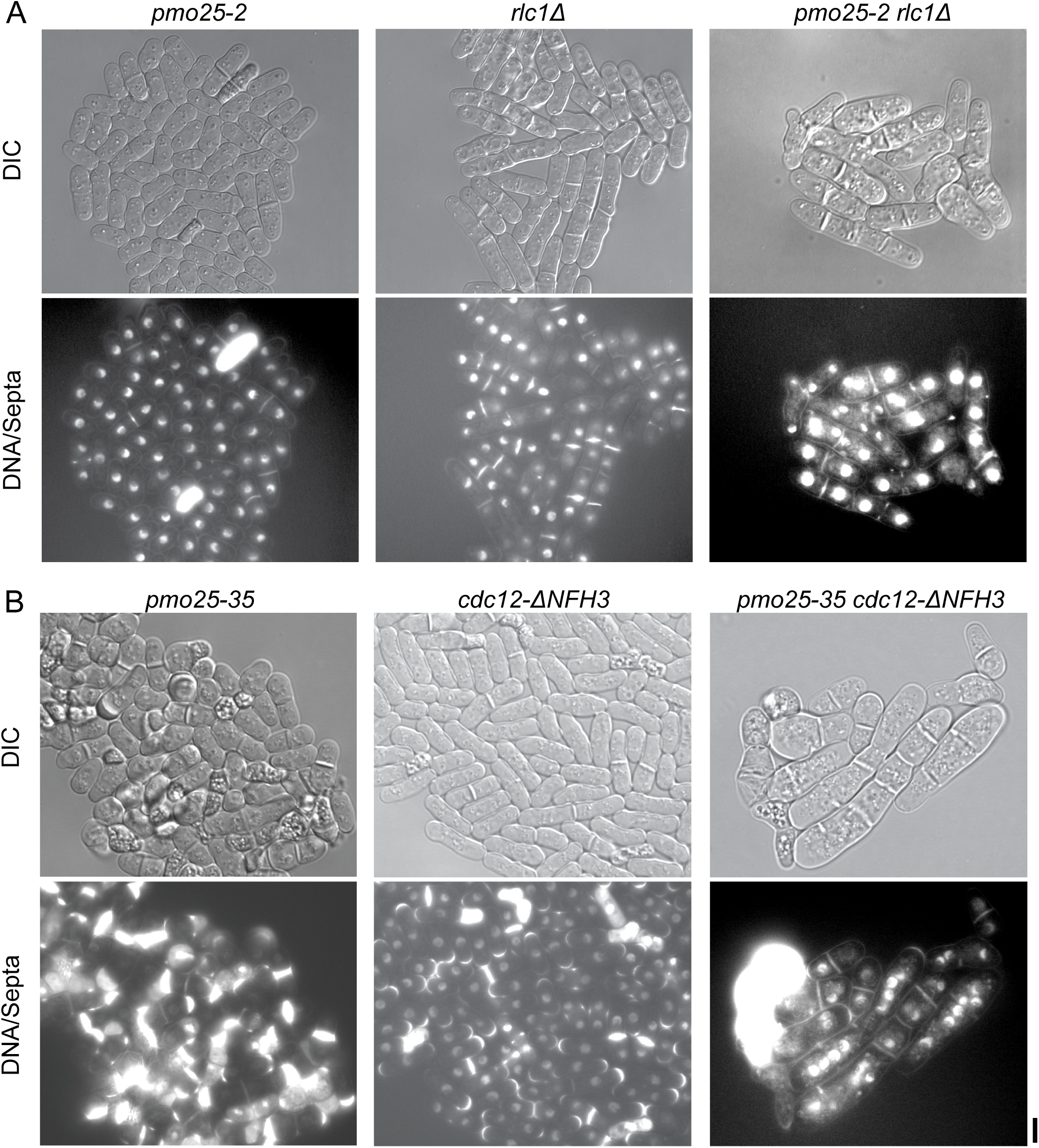
Synthetic genetic interactions between mutations in *pmo25* and the contractile-ring proteins myosin light chain Rlc1 (A) and formin Cdc12 (B). Related to Table 1. Images of DIC and DNA/Septa stained by the Hoechst 33258 (bisbenzimide) dye are shown. Lysed or died cells have the brightest glowing staining that obscures the neighboring cells. (A) Cells were grown at 32°C then shifted to 25°C for 7 h before staining and imaging. (B) Cells were grown at 25°C then shifted to 36°C for 6 h before staining and imaging. Bar, 5 μm.

**Figure S2.**
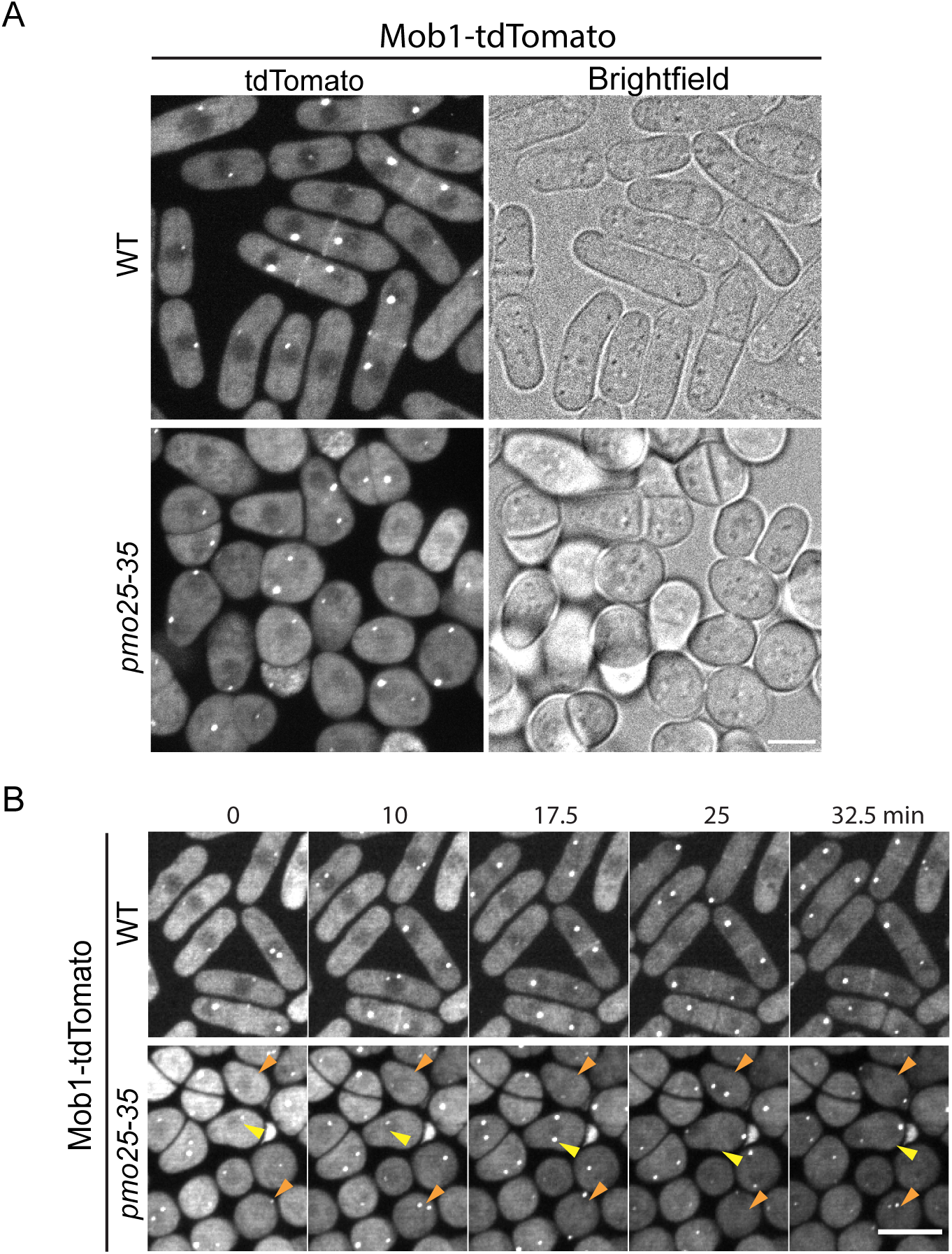
Pmo25 is involved in the recruitment of Mob1 to the division site. Related to Figure 4. (A and B) Single time point images (A) and time course (in min) (B) of Mob1-tdTomato in WT (JW10137) and *pmo25-35* mutant (JW10136) grown at 36°C for 5 h before imaging at 36°C. Arrowheads mark the representative cells with weak or no Mob1 signal at the division site during cytokinesis. Bars, 5 μm.

**Figure S3.**
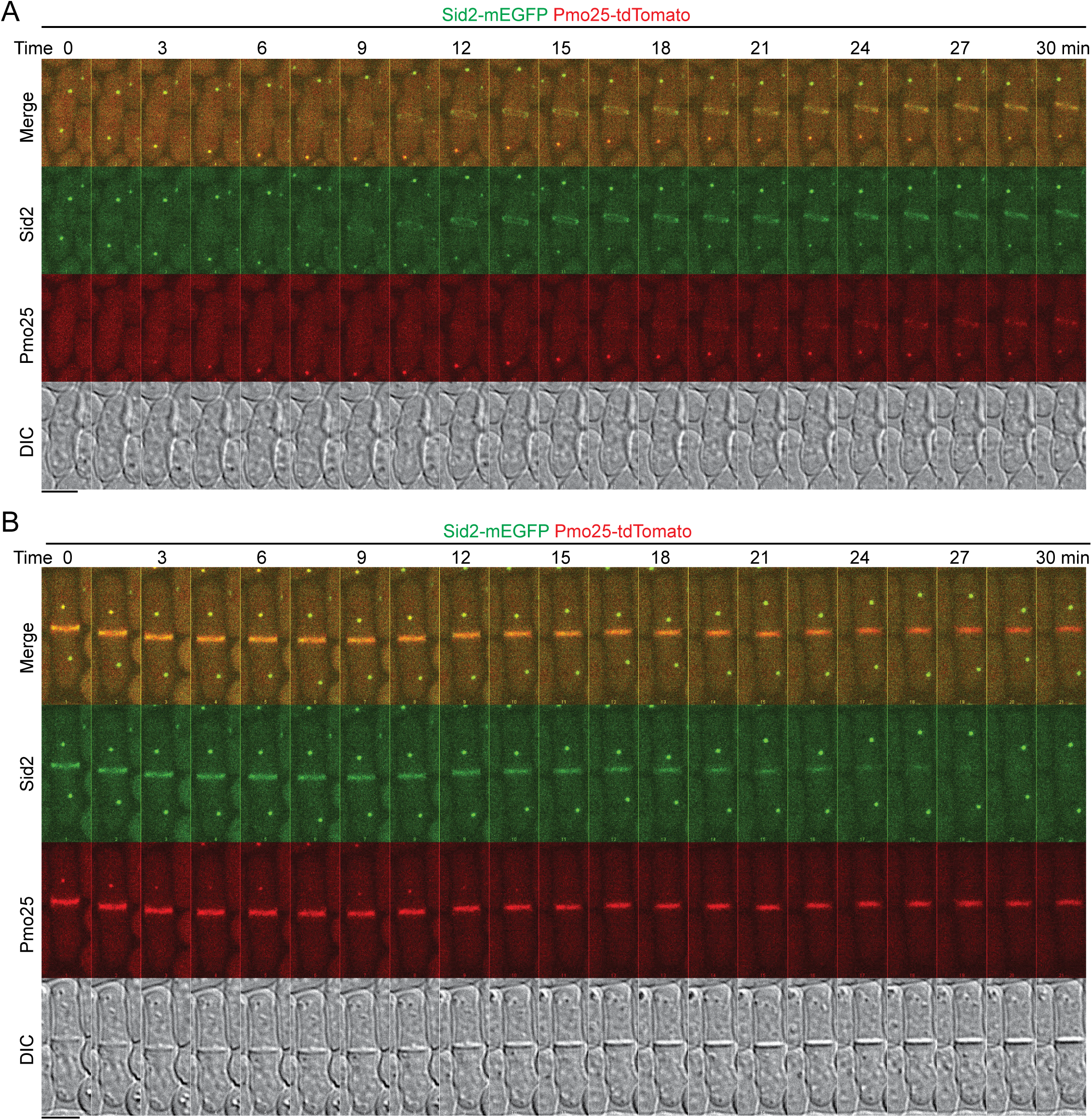
Time courses showing localizations of Sid2 and Pmo25 at the SPBs and the division site during cytokinesis. Related to Figure 5. (A and B) Max intensity projections (9 slices with 0.8 μm spacing) and DIC images showing localizations of Sid2 and Pmo25 in two representative cells. Cells expressing both Sid2-mEGFP and Pmo25-tdTomato (JW10302) were imaged in time-lapse movies over 30 min. Cells were grown at 25°C for ∼48 h before imaging. Bars, 5 μm.

**Figure S4.**
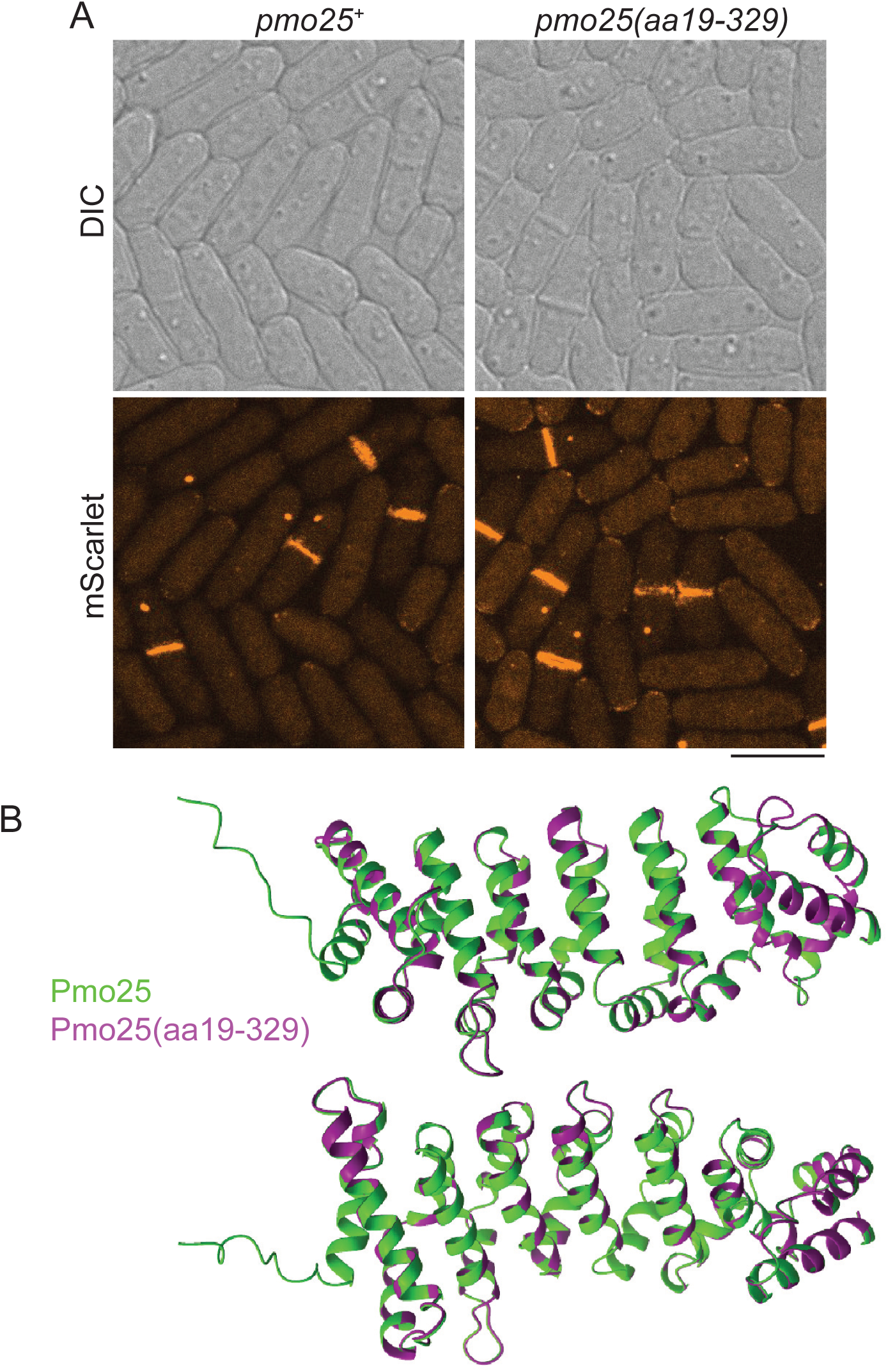
The morphology and localization of Pmo25-mScarlet and Pmo25(aa19-329)-mScarlet NH_2_ terminal truncation mutant. Related to Figure 8. (A) Images of cells expressing Pmo25-mScarlet (JW8034) or Pmo25(aa19-329)-mScarlet mutant (JW8894). mScarlet images are maximal intensity projection of 11 Z slices with 0.5 μm spacing. Bar, 5 μm. (B) Two views of the overlay of the predicted structures of full length Pmo25 (green) and Pmo25(aa19-329) (purple) by AlphaFold3. The models were aligned using the Matchmaker structure analysis tool on ChimeraX.

**Table S1.**
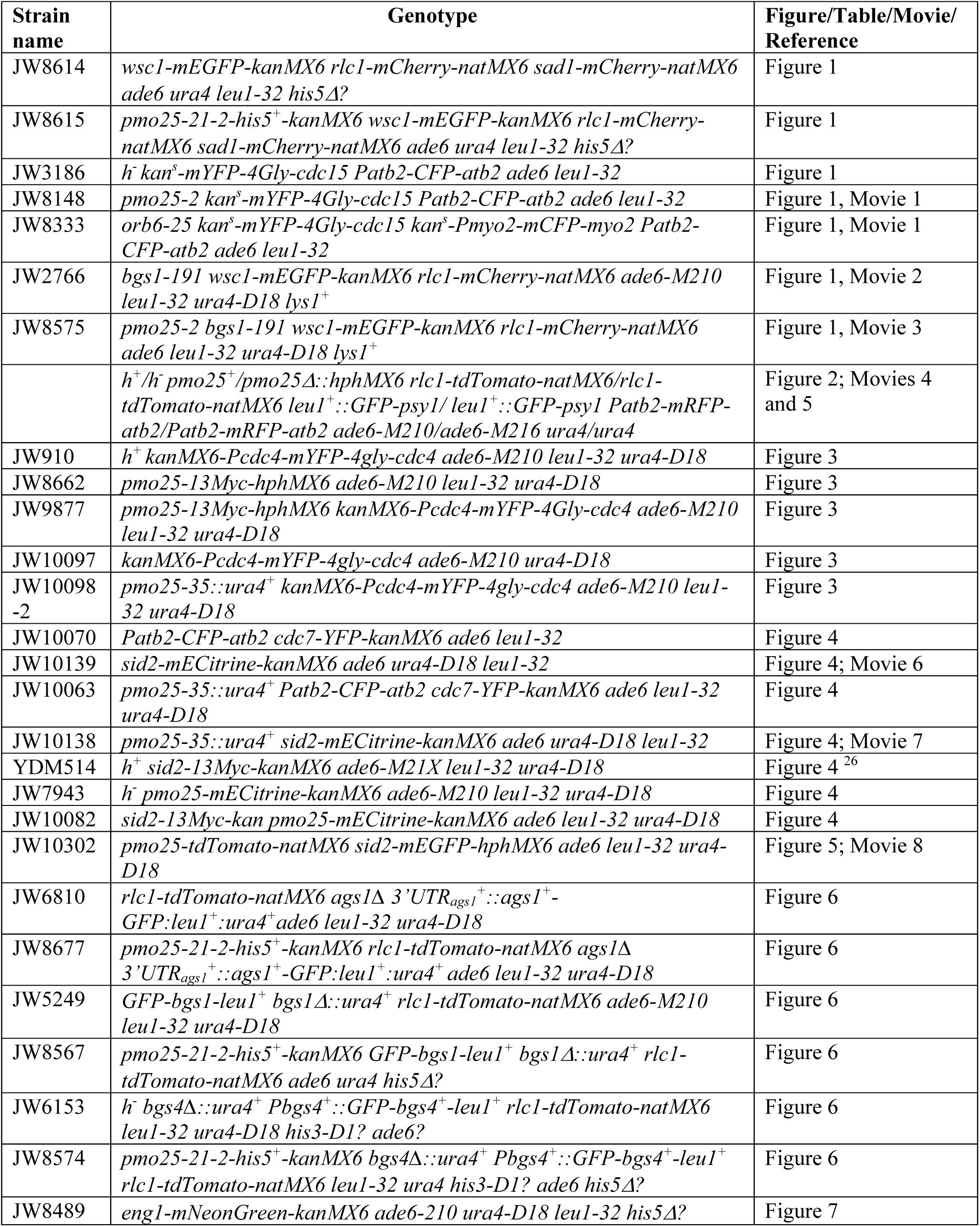
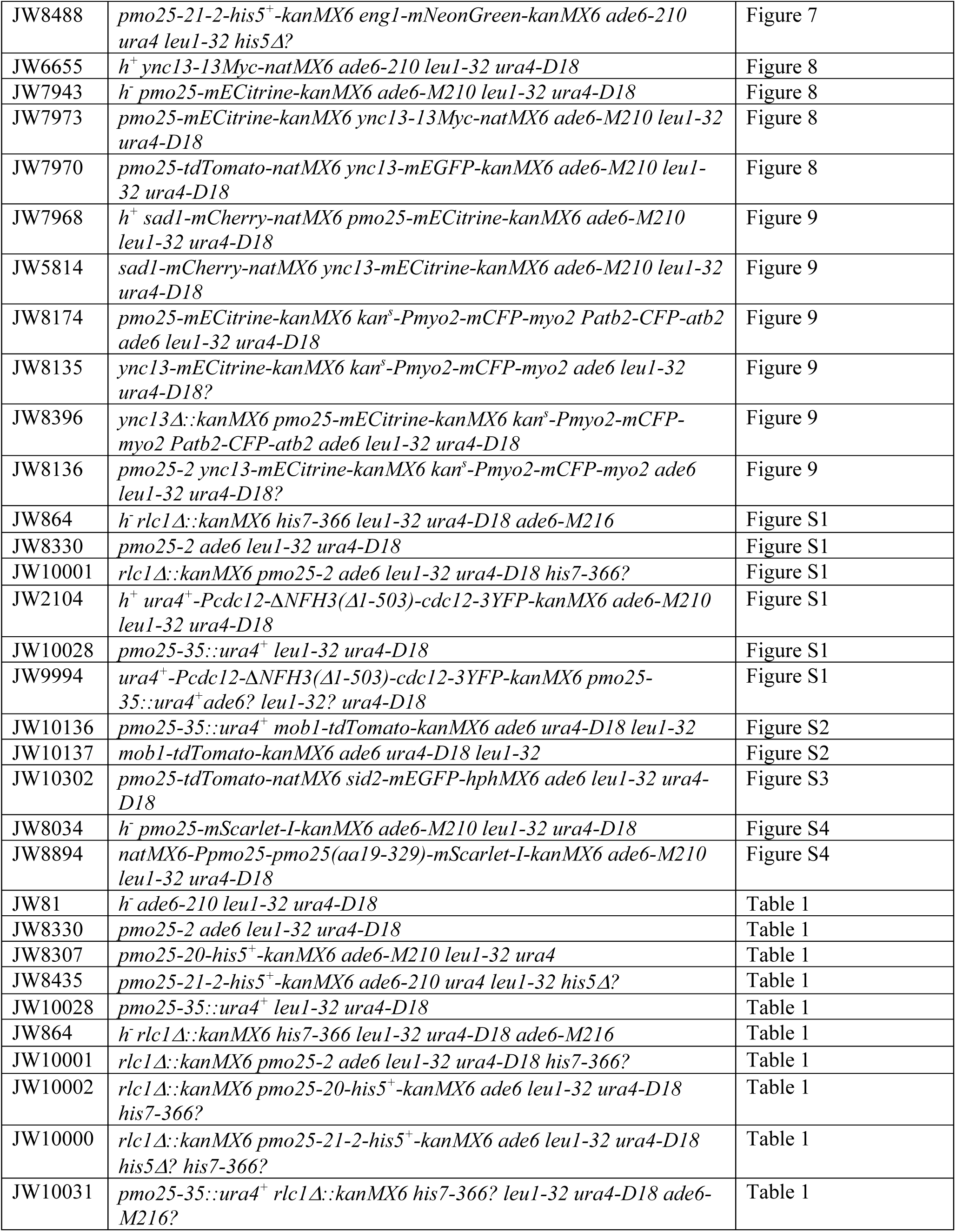
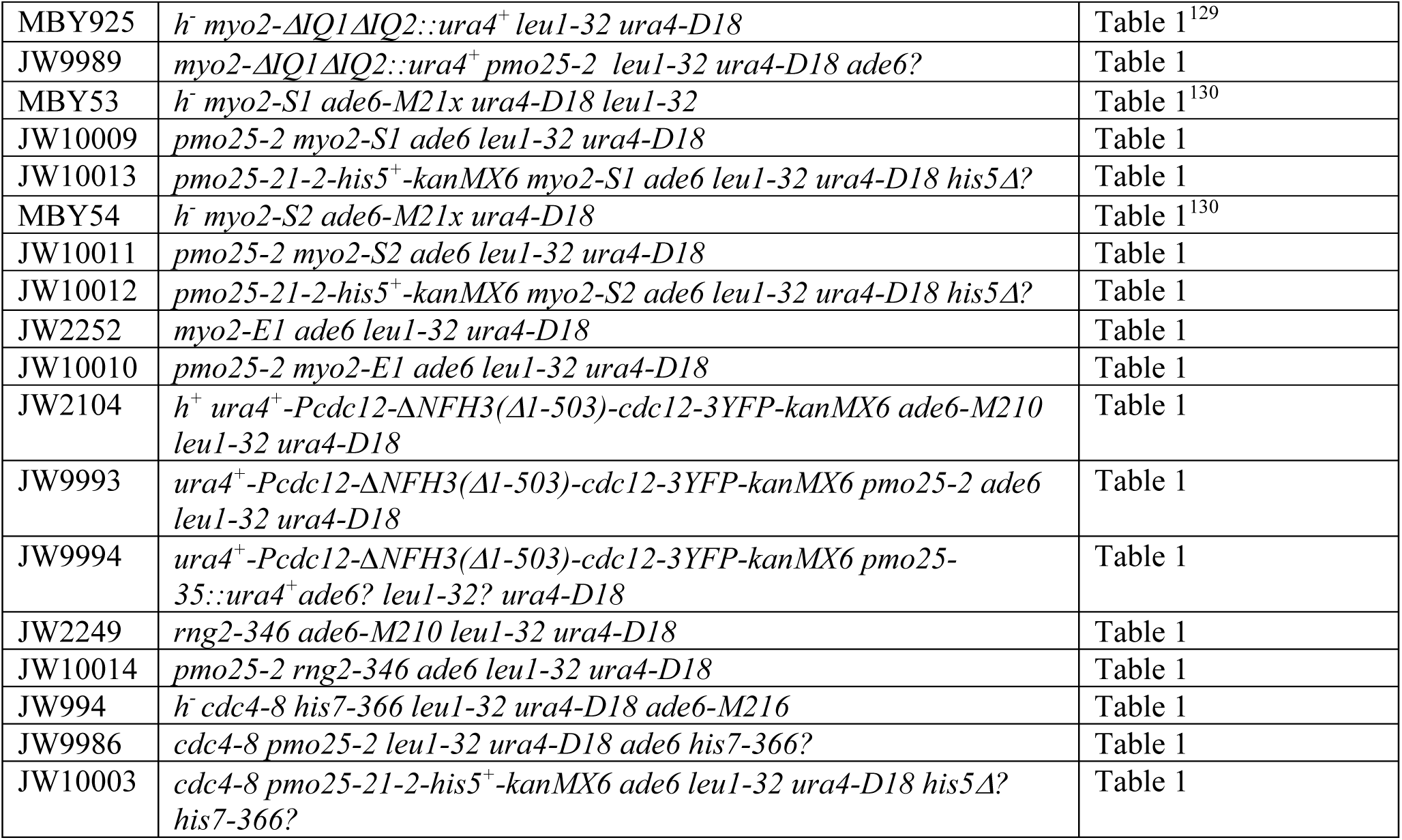
*S. pombe* strains used in this study. Related to the STAR methods.

## Movie legends

**Movie 1. Contractile-ring defects in *orb6-25* and *pmo25-2* cells. Related to Figure 1C**. Cells of *orb6-25 mYFP-cdc15 CFP-atb2 CFP-myo2* (JW8333, left) and *pmo25-2 mYFP-cdc15 CFP-atb2* (JW8148, right) were grown in log phase at 25°C for ∼36 h and then shifted to 36°C for 8 h before imaging at 36°C with 2 min interval. The movie shows the maximal intensity projection of 17 Z slices with 0.5 um spacing. Green: Cdc15; Red: Atb2 Myo2 or Atb2. 7 frames per second (FPS).

**Movie 2. Contractile rings in *bgs1-191* cells. Related to Figures 1D and 1E**. Cells expressing Wsc1-mEGFP Rlc1-mCherry in *bgs1/cps1-191* were grown in log phase at 25°C for ∼36 h and then shifted to 36°C for 2 h before imaging at ∼36°C with 2 min interval. The movie only showed the Rlc1-mCherry channel at maximal intensity projection of 9 Z slices with 0.75 μm spacing. 4 FPS. Bar, 5 μm.

**Movie 3. Contractile rings in *pmo25-2 bgs1-191* cells. Related to Figures 1D and 1E**. Cells expressing Wsc1-mEGFP Rlc1-mCherry in *pmo25-2 bgs1-191* mutant were grown in log phase at 25°C for ∼36 h and then shifted to 36°C for 2 h before imaging at ∼36°C with 2 min interval. The movie only showed the Rlc1-mCherry channel at maximal intensity projection of 9 Z slices with 0.75 μm spacing. 4 FPS. Bar, 5 μm.

**Movie 4. Cytokinesis in *pmo25^+^* cells. Related to Figure 2**. Cells expressing GFP-Psy1 (green) and Rlc1-tdTomato mRFP-Atb2 (red) in *pmo25^+^* WT were grown at 25°C after tetrad dissection as in Figure 2B. Time lapse movie was performed with 8 min interval. Maximal intensity projection of 7 Z slices with 1.0 μm spacing was shown. 2 FPS. Bar, 5 μm.

**Movie 5. Cytokinesis in *pmo25Δ* cells. Related to Figure 2**. Cells expressing GFP-Psy1 (green) and Rlc1-tdTomato mRFP-Atb2 (red) in *pmo25Δ* cells were grown at 25°C after tetrad dissection as in Figure 2B. Time lapse movie was performed with 8 min interval. Maximal intensity projection of 7 Z slices with 1.0 μm spacing was shown. 2 FPS. Bar, 5 μm.

**Movie 6. Sid2-mECitrine localization in WT (*pmo25^+^*) cells. Related to Figures 4B-4E**. Cells expressing Sid2-mECitrine in *pmo25^+^* WT were grown in log phase at 25C°C for ∼36 h, then shifted to 36°C for 4 h before imaging at 36°C. Time lapse movie was performed with 2.5 min interval. Maximal intensity projection of 9 Z slices with 0.75 μm spacing was shown. 1 FPS. Bar, 5 μm.

**Movie 7. Sid2-mECitrine localization in *pmo25-35* cells. Related to Figures 4B-4E**. Cells expressing Sid2-mECitrine in *pmo25-35* were grown in log phase at 25C°C for ∼36 h, then shifted to 36°C for 4 h before imaging at 36°C. Time lapse movie was performed with 2.5 min interval. Maximal intensity projection of 9 Z slices with 0.75 μm spacing was shown. 1 FPS. Bar, 5 μm.

**Movie 8. Sid2-mEGFP and Pmo25-tdTomato colocalization at the SPBs and the division site. Related to Figures 5 and S3.** Cells expressing both Sid2-mEGFP and Pmo25-tdTomato (JW10302) were grown at 25°C for ∼48 h before imaging. Time lapse movies were performed with 1.5 min intervals. Maximal intensity projection of 9 Z slices with 0.8 μm spacing was shown. DIC (top left), GFP (top right), tdTomato (bottom left), merge (bottom right). 2 FPS.

